# MSstatsResponse: Semi-parametric statistical model enhances detection of drug-protein interactions in chemoproteomics experiments

**DOI:** 10.64898/2026.03.09.710598

**Authors:** Sarah Szvetecz, Devon Kohler, Joel D. Federspie, S. Denise Field, Pierre Jean-Beltran, Robert J. Seward, Hyunsuk Suh, Liang Xue, Olga Vitek

## Abstract

Chemoproteomics is a popular approach for the identification of small molecule–protein interactions in biological systems. Several chemoproteomics workflows leverage functionalized chemical probes and mass spectrometry to measure protein engagement through direct protein enrichment or competition using a range of small molecule concentrations. Statistical methods for analysis of such dose-response chemoproteomics datasets are limited. For example, existing methods rely on fixed curve shapes and are sensitive to experimental variation, particularly when the number of doses or replicates is limited. Here, we present MSstatsResponse, a semi-parametric statistical framework for analyzing chemoproteomic dose–response experiments that uses isotonic regression that does not require a fixed curve shape. This approach improves the accuracy and robustness of curve fitting, target identification, and half-response estimation across diverse experimental designs. We evaluate MSstatsResponse by generating a benchmark chemoproteomic dataset that profiled the competition between the kinase-binding probe XO44 and the drug Dasatinib using three mass spectrometry acquisition strategies: data-independent acquisition, tandem mass tag–based data-dependent acquisition, and selected reaction monitoring. We further evaluate the method on simulated datasets that vary the number of doses, number of replicates, and levels of noise, and demonstrate that MSstatsResponse consistently improves sensitivity, specificity, and reproducibility compared to existing methods, particularly in low-replicate and low-dose settings. MSstatsResponse is implemented as an open-source R/Bioconductor package that integrates with the MSstats ecosystem for quantitative proteomics. It provides a unified workflow for preprocessing, curve fitting, target identification, and experimental design, enabling researchers to select the number of doses and replicates appropriate to their experimental goals. The software and documentation are freely available at https://bioconductor.org/packages/MSstatsResponse.

## Introduction

Drug discovery characterizes drug–protein interactions using technologies such as CRISPR-based genetic screens,^1,2^ thermal shift assays,^3,4^ affinity-based profiling,^5^ and cellular imaging.^6,7^ In recent years, chemoproteomics has emerged as a powerful and increasingly popular approach for mapping small molecule–protein interactions directly in biological systems.^8–10^ By integrating functionalized chemical probes with mass spectrometry, chemoproteomics quantifies changes in protein abundance or engagement across a range of small molecule concentration.^8,11–15^ Chemoproteomic experiments have been applied in both basic and translational research to explore drug mechanisms,^16^ identify off-target effects,^17–19^ and prioritize lead compounds.^20^

Experimental goals of mass spectrometry-based chemoproteomic experiments range from exploratory screens profiling targets across many compounds, to confirmatory studies focused on estimating the dose-response curve and on figures of merit such as minimum inhibitory concentration, half effective/inhibitory/occupancy concentration, or toxicity thresholds^11,21–23^ (**Fig. 1**). In competition-based chemoproteomics, these “half-response” quantities are derived from changes in probe-bound protein abundance, and therefore reflect target engagement or occupancy with respect to the assay readout, rather than direct functional inhibition. A major consideration for these experiments is the choice of experimental design, which must balance the range and the number of doses and replicates. While single high-dose screens with biological replicates are common for target identification,^24^ single-replicate, multi-dose designs are favored for broad dose-range profiling.^25^

**Figure 1:**
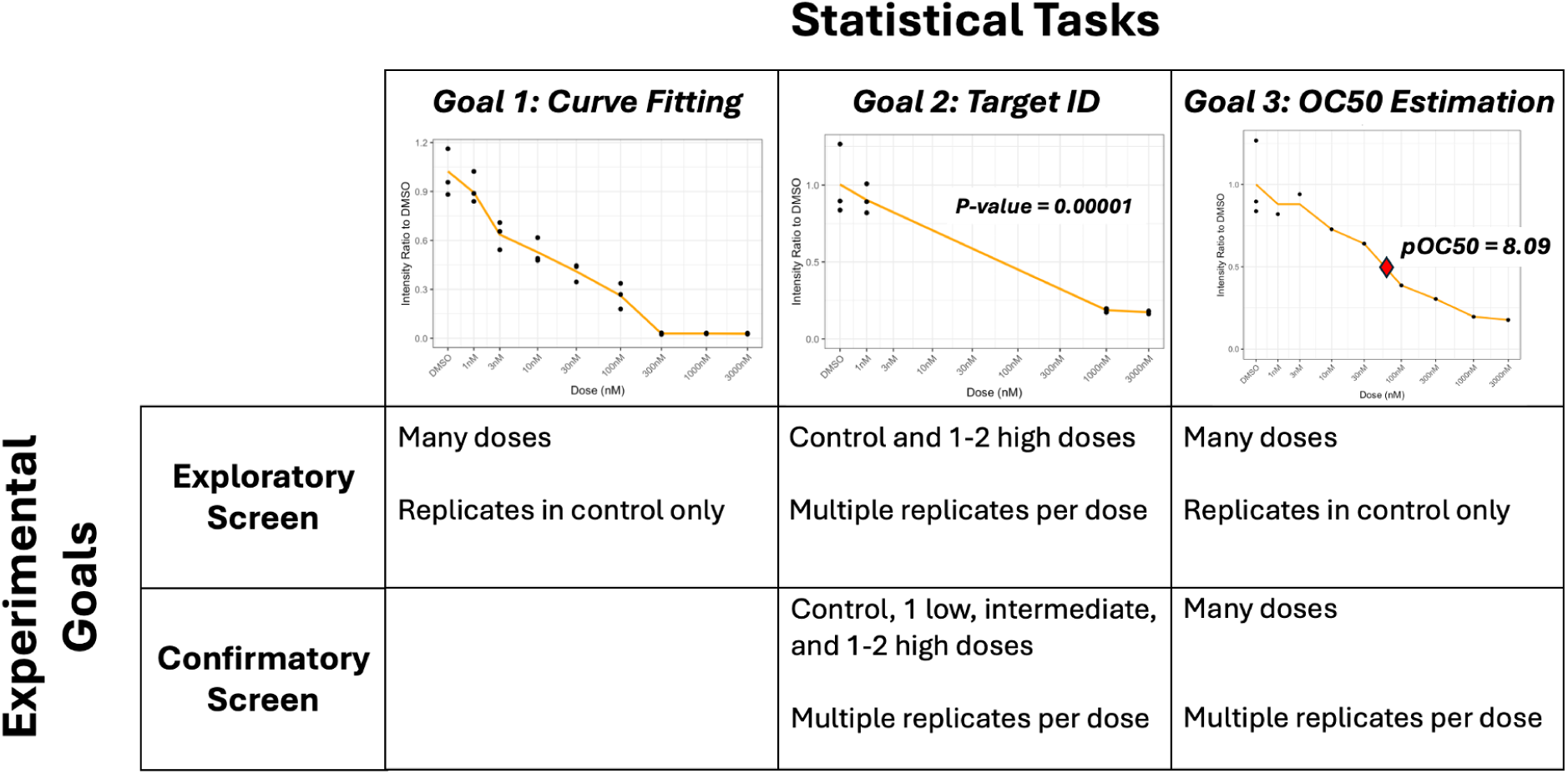
The experimental design of a chemoproteomic experiment depends on the analysis objective. Exploratory screens typically prioritize broad discovery and are optimized for detecting drug–protein interactions using a limited number of doses with more replicates. In contrast, confirmatory screens typically emphasize reproducibility and precision. Accurate OC50 (occupancy) estimation and curve fitting is achieved by including more doses and/or multiple replicates at each dose.

Another consideration is the choice of spectral acquisition. For example, data-independent acquisition (DIA) is often preferred in exploratory settings where the identity of drug-responsive proteins is not known a priori, and broad proteome coverage enables unbiased detection of potential targets and off-targets.^26^ Targeted selected reaction monitoring (SRM), in contrast, is better suited for focused studies on a small, predefined protein set across a limited number of drugs.^27^ Tandem mass tag–based data-dependent acquisition (TMT-DDA) is another alternative for exploratory analyses, often chosen for its ability to reduce missing values within a single multiplexed set, but limited by its multiplexing capacity (typically 18 channels).^28^

Given the diversity of experimental goals, analysis tasks and resource allocations, there is a need for statistical frameworks that support experiments with both exploratory and confirmatory goals and diverse sources of uncertainty and variation, and provide valid statistical inference. Most existing statistical methods fit some form of monotonic curves to the dose–response profile of each protein, and derive figures of merit from these curves. The monotonic nature of the curve is meaningful in the context of chemoproteomics. For example, inhibitors and degraders typically exhibit non-increasing profiles (i.e., decrease in abundance with high dose treatment),^29,30^ while stabilizers and activators produce non-decreasing profiles.^30–32^ The curves often exhibit flat profiles reflecting saturation at both low and high doses, and a slope in the transition zone between the extremes. Therefore, existing curve-fitting packages such as drc,^33^ dr4pl,^34^ and nplr^35^ fit four-parameter log-logistic sigmoid curves and provide parameter estimation and visualization. In many applications this mid-point parameter is reported as an IC50; in this manuscript, for competition-based chemoproteomics we refer to the analogous quantity as OC50 to emphasize that it is derived from probe-bound protein abundance and reflects target engagement/occupancy rather than direct functional inhibition. Despite their usefulness for curve fitting, these approaches lack formal statistical drug–target engagement detection, leaving researchers to manually review large numbers of curves. More recent CurveCurator^21^ addresses this limitation of statistical inference, but relies on the same sigmoidal curve assumption. Such sigmoid curves become unstable in experiments with limited doses, replicates, and high variation, which may result in the loss of weak yet meaningful interactions.^36,37^ Single-replicate experiments are particularly prone to introducing bias, inflating false-positives, and reducing reproducibility of the results.^38^

To address these challenges of statistical analysis we introduce MSstatsResponse, a semi-parametric statistical framework that accounts for the monotonicity of dose–response relationships but does not assume any parametric (e.g., sigmoid) form. Specifically, MSstatsResponse fits isotonic regression to protein-level abundance profiles that is compatible with all experimental goals, experimental designs and mass spectrometry acquisitions, as well as all statistical analysis tasks in **Fig. 1**.

We evaluated the performance of MSstatsResponse by generating a benchmark dataset that profiles the competition between the kinase-binding probe XO44 and the drug Dasatinib using DIA, TMT-DDA, and SRM, as well as on simulated datasets covering a range of experimental designs. Across these evaluations, MSstatsResponse consistently improved sensitivity, specificity, and reproducibility compared to the existing methods, particularly in experiments with limited data. For curve estimation, fewer doses were sufficient to capture meaningful differences in response profiles. For target identification, prioritizing biological replication over dose number improved the sensitivity of detecting drug–protein interactions. For OC50 estimation, incorporating replicates in the control condition and expanding the number of doses improved the precision of the estimation. Based on these results, we developed a practical simulation-based approach to select the appropriate number of doses and replicates in a future experiment for each experimental objective. MSstatsResponse is implemented as an open-source R/Bioconductor package that is freely available as part of the MSstats ecosystem for quantitative proteomics.^39–43^

## Experimental procedures

### Benchmark experiments

A benchmark dataset was generated by pretreated Jurkat cells with varying concentrations of a kinase inhibitor, Dasatinib, followed by treatment of a promiscuous kinase probe, XO44. Briefly, Jurkat cells were grown in RPMI (Thermofisher 11875-093) + 10% FBS (Thermofisher 16000044) + 1X Penicillin/Streptomycin (Thermofisher 15140148). Cells were pretreated with varying concentrations of Dasatinib for 1hr at 37°C in serum free media, followed by treatment of 2 µM XO44 for 30min at 37°C, After probe treatment, cells were spun down and washed with PBS to remove excess probe. Cells were lysed with Phosphate Buffered Saline (14190-144) + 0.25% SDS (L4522)+ 1X HALT PPI (78442) + 1 µL benzonase (712063) followed by sonication (1 sec pulse, 60% amplitude). A biotin handle for enrichment was installed via Click Chemistry (40µM TBTA (T2993), 1mM TCEP (20491), 1mM CuSO4 (451657-10G), 10µM biotin-TAMRA-azide) for 1hr at room temperature. Proteins were precipitated with acetone to remove click reagents, resuspended in PBS + 0.1% SDS, and probe labeled proteins were enriched with high-capacity streptavidin beads (4°C, 16hr). After enrichment, beads were washed with 3x PBS, 3x PBS + 0.1% SDS, 3x water. Beads were resuspended in 2M Urea (ZU10001) + 1mM CaCl2 (21098), proteins reduced with 10mM DTT (D9163-1G), alkylated with 20mM IAA (I11149-5G), and digested with trypsin (37°C, 16hr, V5111). Following digestion, the peptides were desalted (The Nest Group, SNS SS18V) and split into three equal aliquots for the subsequent data acquisition methods.

Positive controls SRC, ABL1, YES, CSK, LCK, and negative control EIF2AK4 were used for benchmarking performance.^44–46^ Response was measured across multiple drug doses and biological triplicates using three acquisition strategies: TMT-DDA, DIA, and SRM. More details on the experimental workflow can be found in **Supplementary Sec. 1**.

The raw mass spectrometry data are publicly available on MassIVE under the accession numbers MSV000100970 (DIA dose-response dataset), MSV000100968 (TMT dose-response datasets), and MSV000100969 (SRM dose-response dataset). The intermediate datasets and analysis scripts used to reproduce the results in this manuscript are available on Zenodo at https://doi.org/10.5281/zenodo.18881723. All benchmark experiments were summarized to the protein level using the same tool (MSstats) to ensure a fair comparison. Uniform processing and normalization steps, handling of poor quality measurements, and treatment of missing values were all the same for every statistical method. A summary of each dataset can be found in **Table 1** and are described in further detail below.

**Table 1:** Data overview and availability. “No. of doses” is the number of drug doses (concentrations) in each dataset. “No. of replicates” is the number of replicates per drug dose; simulations were generated with varying replicate numbers. “No. of proteins” is the total number of proteins quantified. “Average no. of peptides” is the average number of peptide measured per protein. “Positive controls” are the five known Dasatinib targets (SRC, ABL1, YES, LCK, and CSK), and “Negative controls” is EIF2AK4. “Data availability” refers to the MassIVE repository ID for experimental datasets or the GitHub repository for simulations.

| **Metric** | **DIA** | **TMT** | **TMT-RTS** | **SRM** | **Simulation** |
| --- | --- | --- | --- | --- | --- |
| No. of doses | 9 | 6 | 6 | 9 | 2/4/5/9 |
| No. of replicates | 3 | 3 | 3 | 3 | 1/2/3/4 |
| No. of proteins | 7,329 | 5,935 | 6,668 | 175 | 3,000 |
| Average no. of peptides | 17 | 8 | 8.7 | 1 | N/A |
| Positive controls | 5 | 5 | 5 | 5 | 2,000 |
| Negative controls | 1 | 1 | 1 | 0 | 1,000 |
| Data availability | MASSIVE | MASSIVE | MASSIVE | MASSIVE | Github |

**Dataset 1: DIA Benchmark** Samples were run on a Bruker timsTOF SCP using default acquisition parameters. The DIA dataset comprises 27 samples, including triplicate DMSO controls and eight drug doses (1 nM, 3 nM, 10 nM, 30 nM, 100 nM, 300 nM, 1 µM, and 3 µM). Data were processed using Spectronaut with default parameters.

**Dataset 2: TMT Benchmark** The desalted peptides were dried down and resuspended in 100 mM EPPS, pH8.5 (ThermoFisher Scientific, J61476.AE) at a final concentration of 30% acetonitrile and were reacted with 125 µg of TMTpro reagent for 1 hr at room temperature. The TMT reaction was quenched with the addition of ammonium hydroxide to a final 0.5% concentration. The samples were then pooled and desalted and the TMT labeled peptides were fractionated using the Pierce High pH Reversed-Phase Peptide Fractionation Kit (ThermoFisher Scientific 84868) according to manufacturer’s instructions. LC-MS^3^ analysis of the TMT samples was performed on a Dionex Ultimate 3000 UPLC system (ThermoFisher Scientific, CA), coupled online to an EASYSpray ion source and an Eclipse Tribrid mass spectrometer with FAIMS (ThermoFisher Scientific, CA). TMT-labeled peptide fractions were separated on an EASYSpray C18 column (75 µm × 50 cm, ES903) heated to 50°C using mobile phases A (0.1% formic acid in water) and B (0.1% formic acid in 90% acetonitrile) at a flow rate of 250 nL/min with a linear gradient from 8% B to 28% B for 117 min followed by a ramp to 35% B over 10 min. The real time search (RTS) data was acquired using previously described instrument settings.^47^ The SPS MS3 dataset was acquired using identical instrument settings but without out the real time search step.

**Dataset 3: SRM Benchmark** Selected Reaction Monitoring (SRM) was performed using a TSQ Altis Mass Spectrometer with an EASY-Spray Ion Source (ThermoFisher Scientific) coupled with a Dionex UltiMate 3000 UHPLC (ThermoFisher Scientific). Peptides were first trapped on a C18 trap cartridge (#174500, ThermoFisher Scientific) and washed with 0.5% trifluoroacetic acid for 2 minutes. They were then separated on an EASY-Spray column (#ES900, ThermoFisher Scientific) at a flow rate of 400 nL/min over a linear gradient: 1-45% B, 2’-25’ (solvent A: 2% acetonitrile 0.1% formic acid, solvent B: 90% acetonitrile 0.1% formic acid). Subsequently, the trap cartridge and analytical column were each washed with 80% acetonitrile 0.1% formic acid and equilibrated in either 0.5% trifluoroacetic acid or 1% A, respectively, prior to the next injection. The column temperature was maintained at 50°C. Peptides were ionized at a spray voltage of 2.8 kV in positive ion mode while the ion transfer tube temperature was maintained at 300°C. SRM was carried out on 175 proteins, using one peptide per protein and two precursor ions for each peptide (one light, one heavy). Full SRM table can be found in Supplemental Information (Supplemental Table S1). Key parameters included scheduled acquisition with a 90 sec retention time window, a cycle time of 2 sec, Q1 resolution (FWHM) of 0.7, and Q3 resolution (FWHM) of 1.2. After data was acquired, peak area integration was conducted using Skyline. The SRM dataset includes 27 samples, consisting of triplicate DMSO controls and the same eight drug doses as the DIA dataset (1 nM to 3 µM).

### Simulated datasets

To evaluate method performance under different experimental designs and conditions, we simulated chemoproteomic datasets based on template protein abundances from the DIA benchmark (**Supplementary Fig. 3**). Template proteins were defined as either strong or weak interactors (true positives) or as non-interactors (true negatives). Strong drug–protein interactions corresponded to proteins whose probe-bound abundance decreased to near zero at the highest drug dose, reflecting maximal competition (i.e., near-complete target engagement by the free drug). Weak drug–protein interactions represented proteins whose probe-bound abundance decreased to approximately 50% of their baseline level, reflecting partial engagement. True negatives maintained constant abundance across all doses, indicating no drug effect. Specifically, the SRC protein served as a strong interactor, TEC as a weak interactor, and EIF2AK4 as a non-interactor. We then generated multiple experimental scenarios by varying the number of doses, biological replicates, and amount of variation to benchmark each method across a range of design settings. Full simulation details are provided in the **Supplementary Sec. 1.2** and code on Zenodo at https://doi.org/10.5281/zenodo.18881723.

## Background

### Input data and notation

Statistical analyses of chemoproteomic experiments typically take as input protein-level abundance values. For acquisitions such as DIA, DDA, SRM or PRM, the spectral features are first processed with pipelines such as Skyline,^48^ Spectronaut,^49^ FragPipe,^50^ Proteome Discoverer^51^ or MaxQuant^52^ that detect, identify and quantify spectral features and summarize the feature abundances into a single value per protein per run. Alternatively, specialized packages such as MSstats,^40^ QFeatures,^53^ MaxLFQ,^54^ or MSnbase^55^ take as input the identified and quantified spectra features and provide dedicated statistical frameworks for protein summarization and quality control. Important aspects of protein-level summarization include normalization of feature or protein abundances between MS runs and treatment of missing values.

### Existing statistical methods for dose-response analysis

In this manuscript, we consider a protein measured in an experiment with *r* = 1*, …, R* biological replicates at each of the *d* = 0, 1*, …, D* doses, where *d* = 0 denotes a control such as DMSO. Therefore, the total number of measurements is *R*(*D* + 1). Let *x_d_* denote the drug dose at index *d*, and *a_dr_* denote the corresponding protein abundance in replicate *r* on the original (i.e., not log-transformed) scale.

The most popular model for dose-response analysis in chemoproteomics is the four-parameter log-logistic model (4PL), also known as the Hill model, implemented in tools such as drc,^33^ dr4pl,^34^ and CurveCurator.^21^ These models assume a sigmoidal response as summarized in **Table 2**. From the curve fit, the tools estimate the lower and upper asymptotes, slope (Hill coefficient), and OC50 parameters as detailed in **Supplementary Table 1**. **drc** This R package fits and visualizes dose-response curves. It is not specifically designed for chemoproteomics experiments, but can easily be applied to any dataset that quantifies dose–response relationships. Prior to curve fitting, users must transform the input protein abundances at each dose and replicate relative to the control, i.e.

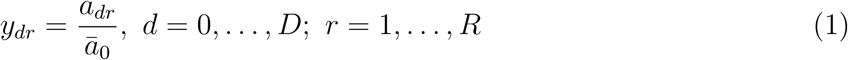

**Table 2:** Curve fitting models and input protein abundance scales of the existing and proposed approaches.

| Method | Input | Model |
| --- | --- | --- |
| **drc** | Ratio to DMSO (**Eq. 1**); original scale | $y_{dr} = \theta_1 + \frac{\theta_4 - \theta_1}{1 + 10^{\theta_3(x_d - \log_{10} \theta_2)}} + \epsilon_{dr}$ $\epsilon_{dr} \stackrel{\text{iid}}{\sim} N(0, \sigma^2)$ |
| **dr4pl** |  |  |
| **CurveCurator** | Ratio either to DMSO (**Eq. 1**) or to abundance at $\min(x_d)$; original scale | |
| **ANOVA** | $\log_2$ scale | $y'_{dr} = \mu + x_d + \epsilon_{dr}$ $\sum_{d=0}^D x_d = 0; \epsilon_{dr} \stackrel{\text{iid}}{\sim} N(0, \sigma^2)$ |
| **MSstatsResponse** | *Curve fitting and OC50 estimation:*<br>ratio to DMSO (**Eq. 1**); original scale. *Target identification:*<br>$\log_2$ scale | $y'_{dr} = f(x_d) + \epsilon_{dr}$ $\epsilon_{dr} \stackrel{\text{iid}}{\sim} N(0, \sigma^2) \text{ where } f(x_0) > f(x_1) > \dots > f(x_D)$ |

where *ā*_0_ is the mean protein abundance in the control. This transformation rescales the data such that in a four-parameter log-logistic model the response is expressed as a percent reduction relative to the control (i.e., 0–100%). Parameters of the curve are then estimated by minimizing the sum of squared residuals. For each of the four model parameters (i.e., lower and upper asymptote, slope, and mid-point concentration) the package reports the estimated values along with the fitted sigmoid curve.

While drc addresses curve-fitting and OC50 estimation tasks, it does not provide a formal hypothesis testing framework for identifying significant drug–protein interactions. The package provides standard errors and p-values for each of the four model parameters, and tests the null hypothesis of each parameter equal to a certain value (e.g. slope = 0), however it does not assess the overall curve fit. As a result, many users choose to manually inspect pages of dose-response curves, or apply filtering based on slope and OC50 estimate specific p-values, to find drug-protein interactions.

#### dr4pl

Developed as an alternative to drc, this R package addresses several of its limitations (in particular, model fitting convergence issues) to provide a more robust framework. Like drc, it is not designed specifically for chemoproteomics experiments, but can be applied with similar data preparation. In practice, it takes as input protein abundances transformed as in **Eq. 1**, and fits and visualizes a four-parameter log-logistic curve to estimate the mid-point concentration. Like drc, dr4pl returns 95% confidence intervals for all model parameters (**Supplementary Table 1**). However, this R package does not improve on the hypothesis testing task for identifying significant drug–protein interactions. Users must define their own criteria for identifying interacting proteins based on model outputs, such as checking whether the confidence interval for the slope contains zero, or whether the confidence interval for OC50 lies within the experimental dose range.

#### CurveCurator

This Python package was developed specifically for chemoproteomics-like experiments and integrates normalization, curve fitting, and statistical inference into a single workflow. Unlike drc and dr4pl, it takes as input summarized protein abundances, *a_dr_*. Within the workflow, it automatically applies the transformation in **Eq. 1**, and provides alternative transformations relative to the smallest drug dose (or another chosen control). Upon the transformation, it facilitates optional steps such as global median centering, imputation of missing values using intensity-based rules (e.g., imputing low-intensity values from a distribution), and filtering out curves with excessive number of missing values.

CurveCurator fits the same four-parameter log-logistic model as drc and dr4pl and pro-vides single-point OC50 estimates. In contrast to the other methods, it implements a formal hypothesis test for drug–protein interaction detection by comparing each fitted curve to a null model of constant response with a recalibrated F-statistic. This recalibration uses simulated null curves to estimate effective degrees of freedom and p-values. Beyond the p-values, CurveCurator combines statistical significance with effect size (log2 fold-change between lowest and highest dose) into a single relevance score. Curves are then classified as significantly up-, down-, not regulated, or unclear (for high-variance cases). The developers recommend including more drug doses rather than additional replicates at each dose level.^56^

### Existing general statistical methods for proteomic experiments

All the packages above transform the input protein abundances at each dose and replicate relative to a control. While fitting the curve on this scale helps visual assessment and enables direct estimation of OC50, this transformation has negative consequences. For example, if the variability in the DMSO condition is not properly estimated (e.g., due to lack of replicates), the resulting curves shapes may become distorted. Alternatively, if variability is high within the control replicates, then that noise is propagated through all other measurements. Therefore, it is attractive to consider alternative statistical analyses that do not require such transformation.

#### ANOVA

Analysis of variance (ANOVA) is a general statistical approach implemented in most programming languages. It is not specifically designed for proteomics or dose–response experiments, but is easily applied. In the context of proteomics, it translates to a test of differential protein abundance across dose conditions. Similarly to drc, dr4pl and CurveCurator it takes as input protein abundance values, however the ratio-scaled values are not required. Instead, ANOVA compares the means of log_2_-transformed protein abundances at each *D* + 1 doses

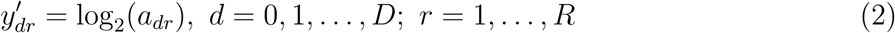

ANOVA requires at least two replicates for at least one dose, however designs with replicates at each dose perform best. The ANOVA model assumes that each dose has its own expected value of protein abundance and an equal variance of replicates for each dose, including for the control such as DMSO (**Table 2**). Parameters of the model are estimated using least squares. Hypothesis test compares the means of protein abundance at each dose to the null curve with a constant response using F test (**Supplementary Table 1**).

Since ANOVA avoids taking ratios of protein abundances relative to a control, it is not susceptible to the artifacts of this approach. Moreover, ANOVA does not assume any functional relationship between the doses and the response, providing additional flexibility. However, this flexibility has its own drawbacks. In particular, users have to manually format protein abundances, manually perform normalization and missing value imputation if needed, and interpret the ANOVA output. Furthermore, the fitted profiles can exhibit non-monotonic hook-like patterns that may reflect biologically implausible artifacts. The assumption of constant variance between DMSO and drug-treated protein abundances may not hold. Finally, ANOVA does not naturally support the estimation of OC50.

#### MSstats

MSstats is a family of R packages developed for statistical analysis of mass spec-trometry–based proteomics data.^40,57^ MSstats supports input from all major mass spectrometry acquisitions such as DDA and TMT-DDA, DIA and SRM, and includes converters for many widely used spectral processing tools such as Skyline,^48^ Spectronaut,^49^ FragPipe,^50^ Proteome Discoverer,^51^ and MaxQuant^52^ that identify and quantify spectral features. The processing workflow supports steps for quality control, multiple normalization strategies (e.g., median, reference-based, global), optional imputation for missing values using an accelerated failure time (AFT) model, and protein summarization from peptide or fragment-level data. Therefore, MSstats can be applied to datasets generated from a range of experimental questions and pipelines, and supports the reproducibility of downstream statistical analyses.^57^

MSstats log-transforms feature intensities prior to summarization, and therefore protein-level summaries are comparable to the log_2_ scale. Statistical analyses in MSstats model these protein-level abundances, and extend ANOVA using a flexible family of linear mixed-effects models applicable to complex designs. In dose–response experiments, MSstats supports comparisons of specific doses (e.g., DMSO versus the maximum dose) using user-defined contrasts.^58^ Although MSstats is better suited for chemoproteomic experiments than ANOVA by automating the workflow and providing more flexible models, the available statistical analyses have the same drawbacks as ANOVA and currently do not support all the tasks necessary for chemoproteomics.

## Results

### Overall workflow in MSstatsResponse

The proposed approach implemented in MSstatsResponse is overviewed in **Fig. 2**. The MSstatsResponse is implemented in an open-source R package fully integrated with the MSstats ecosystem, available at https://bioconductor.org/packages/MSstatsResponse.

**Figure 2:**
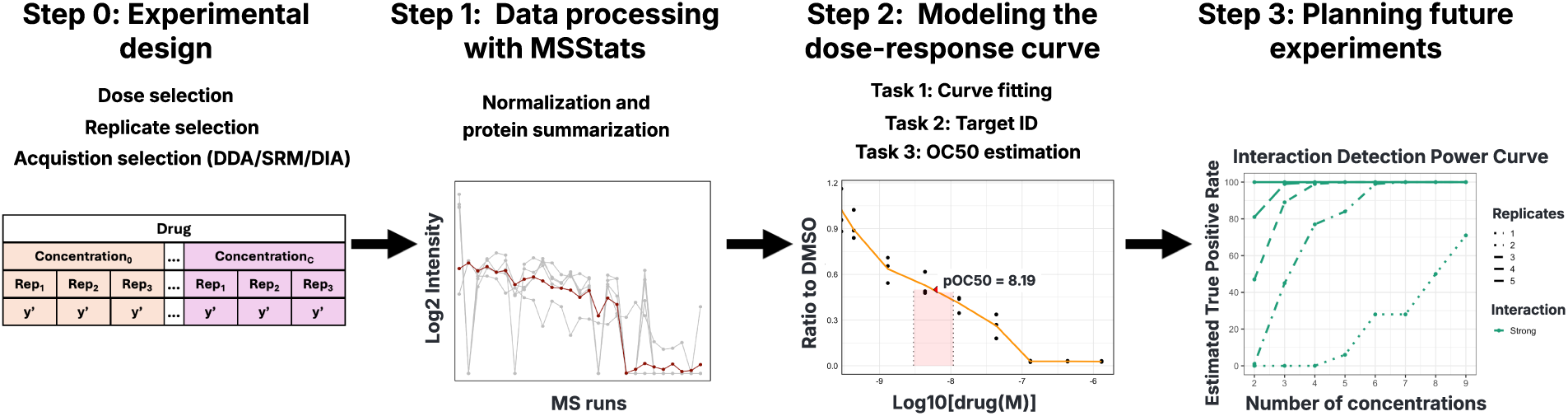
Proposed workflow for chemoproteomic experiment data analysis with MSstatsResponse. Step 0: the workflow starts with the experimental design, which includes selection of doses, replicates and acquisition. Step 1: the acquired data are normalized, and protein abundances are summarized using MSstats. Step 2: MSstatsResponse supports three analysis tasks: curve fitting, target identification, and OC50 estimation. Step 3: the existing dataset serves as a basis for selecting appropriate number of doses and replicates in future investigations.

#### Step 0: Experimental design

Since different analysis goals require different trade-offs between the number of drug doses and biological replicates (exploratory screens often prioritize more doses with fewer replicates, while confirmatory studies may emphasize replication with fewer doses), MSstatsResponse supports all such designs. MSstatsResponse also supports unreplicated experiments for curve fitting and OC50 estimation, however evaluating the statistical significance of target identification requires replicates in at least one dose.

#### Step 1: Data processing with MSstats

To optimize this step for quantitative proteomics, MSstatsResponse builds directly on the MSstats infrastructure for preprocessing, normalization, and protein summarization. MSstats provides the option of global median normalization, appropriate when the drug is expected to affect only a small number of proteins. If the drug is expected to affect many proteins, median normalization should be turned off, and the data should be normalized with respect to reference protein standards or be left without normalization. MSstats excludes proteins with an excessive number of missing values and imputes the remaining values using an accelerated failure time model.^59,60^ The resulting protein-level summaries serve as input for all subsequent MSstatsResponse analyses. However, using MSstats is not required, and the remaining steps are in principle compatible with other appropriate data processing and protein summarization methods and software.

#### Step 2: Modeling the dose-response curve

This step is the main contribution of this manuscript. Similarly to ANOVA and MSstats, we propose to analyze protein-level summaries on the log-transformed scale (as opposed to the ratio-transformed scale). Similarly to drc, dr4pl and CurveCurator, MSstatsResponse includes a monotonic relationship constraint between the dose and the response to avoid overfitting to experimental artifacts. However, the monotonic relationship is not restricted to a strict sigmoid shape.

To fit such a curve, MSstatsResponse relies on isotonic regression (**Table 2**).^61,62^ Isotonic regression is a non-parametric statistical model that specifies a piecewise linear function under a monotonicity constraint. Its utility has been demonstrated in dose-finding in clinical trials,^63^ high-throughput genotoxicity screening,^64^ and causal dose–response estimation.^65^ In these applications, isotonic regression was favored for its greater flexibility as compared to parametric curves, and its robustness in fitting non-sigmoidal curve shapes. However, its utility for chemoproteomics has not been explored so far. MSstatsResponse relies on the robustness of isotonic regression to irregular or noisy patterns and on its ability to easily accommodate various experimental designs and analysis tasks. The three primary analysis tasks (curve fitting, target identification, and OC50 estimation) are described in more detail below.

#### Task 1: Curve fitting

By default, MSstatsResponse specifies a non-increasing constraint, reflecting the expected trend in inhibitor screens, but this can be easily reversed to model non-decreasing responses when appropriate. Under the non-increasing constraint, the method minimizes the weighted squared error between the observed protein abundances and a function *f* (*x_d_*)^61,62^

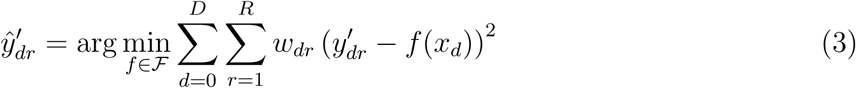

where 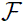 is the set of all piecewise linear functions that satisfy the constraint, and *w_dr_* are optional observation weights.

Parameters of *f* (*x_d_*) are estimated using the Pool-Adjacent-Violators Algorithm (PAVA).^61,62^ In an unreplicated experiment (*R* = 1), PAVA begins with the observed abundances at each dose *d*, *y*^′^_*d*1_, as the fitted values. It then scans through adjacent dose levels and checks whether the monotonicity constraint is violated, i.e. whether 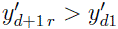, to ensure a non-increasing fit. If the monotonicity constraint is violated, PAVA iteratively pools adjacent dose levels until the constraint is satisfied. This process continues until all adjacent pairs satisfy the monotonicity constraint. The resulting fit is a piecewise linear, non-increasing function. In replicated experiments (*R >* 1), the initial fitted values are replaced by the mean abundances at each dose 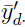. The default in MSstatsResponse is set to equal weights. In experiments with unequal weights, pooled values are replaced with their weighted averages. This approach is useful, e.g., in experiments with unequal quality measurements.^66^

Isotonic regression is compatible with both log-transformed and ratioed measurements. Therefore, MSstatsResponse uses different scales depending on the analysis task. For curve fitting and visualization, MSstatsResponse models the data on the ratioed scale, since this aligns with the familiar convention of expressing dose–response relationships as a percent change relative to control. In competition-based chemoproteomics this corresponds to the fraction of probe-bound protein remaining as a function of drug concentration. Since protein-level abundances reported by MSstats are comparable to the log_2_ scale, the task begins by exponentiating the summaries to recover the original-scale intensities, i.e. 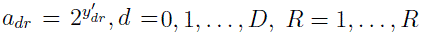. These values are then normalized relative to the control condition as in **Eq. 1**, yielding the ratio-scale responses *y_dr_*. In all cases, the *x*-axis (drug dose) is transformed to log_10_ molar units, following the guidance in.^36^

#### Task 2: Target identification

For target identification, MSstatsResponse fits the isotonic regression on the log-transformed (as opposed to ratioed) scale to better match the assumptions of the downstream statistical inference. To detect drug–protein interactions, MSstatsResponse tests the null hypothesis of constant protein abundance across all doses, i.e. 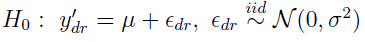 versus the alternative 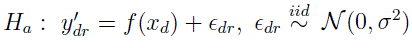 where *f*(*x*) is as in **Eq. 3** and *d* = 0, 1, …,*D*, *r* = 1, …,*R*, using standard F-test (**Supplementary Table 1**). P-values derived from the F-distribution are adjusted for multiple testing using the Benjamini–Hochberg procedure to control the False Discovery Rate.

The log transformation 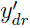 for this step was chosen based on methodological considerations, specifically on the fact that dividing all the observed values by a common observed reference violates the assumption of independence of the transformed values. The choice was also confirmed empirically through simulations. **Supplementary Fig. 5** illustrates that the log scale increased the sensitivity of detecting true interactions without inflating the false positives as compared to the ratioed scale, notably for weak interactions.

#### Task 3: OC50 estimation

In this manuscript we define OC50 as the drug concentration at which the probe-bound protein abundance is reduced to 50% of its control level. In a competition-based chemoproteomics assay, this corresponds to the point where half of the probe-binding capacity has been competed away by the free drug, and therefore reflects target engagement or occupancy with respect to the assay readout rather than direct functional inhibition. Since the curves on the ratioed scale have an intuitive interpretation as percent change relative to control,^36^ MSstatsResponse estimates OC50 on the ratioed scale as in Task 1.

Since the isotonic regression curve *f* (*x_d_*) is piecewise linear, MSstatsResponse linearly interpolates between the values *f* (*x_d_*) *>* 0.5 and *f* (*x_d_*_+1_) *<* 0.5 that are closest to 0.5

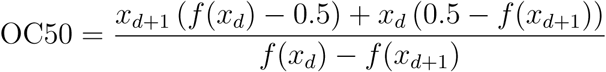

The parameter is estimated using fitted values *f*^^^(*x_d_*). If all *f*^^^(*x_d_*) *>* 0.5, the OC50 is labeled as undefined. A similar approach can estimate other figures of merit such as OC25, OC95, etc.

All OC50 values in MSstatsResponse are reported as pOC50 (i.e., -log_10_OC50) values. This follows standard reporting practice with the field,^36^ where larger values indicate higher potency. For simplicity, we refer to pOC50 values as OC50 throughout the text.

Confidence intervals associated with the estimated OC50 are obtained using stratified bootstrap resampling (default *n* = 1000). To preserve the replicate structure of the experiment the procedure resamples the observations separately within each dose, estimates OC50 from each resampled dataset, and reports the quantiles of the empirical distribution of resampled estimates. Non-stratified bootstrapping is used for single-replicate datasets.

Although MSstatsResponse estimates OC50 on the ratio-transformed scale, the approach is in principle compatible with the log scale as described in **Supplementary Table 1**. **Supplementary Fig. 6** illustrates that the resulting estimates and confidence intervals are similar on both scales.

#### Step 3: Planning future experiments

Statistical experimental design evaluates the performance of a future experiment with various combinations of doses and replicates, and given realistic estimates of systematic and random biological and technological variation. MSstatsResponse implements functionalities for such experimental design, treating the current dataset as representative of future investigations.

First, MSstatsResponse creates systematic dose-response profiles representative of future strong, weak, and non-interacting proteins. Users specify the unique identifiers (e.g., protein IDs) corresponding to these representative proteins, which are selected from the fitted dose–response curves in the current dataset. The tool can use one, two, or all three interaction types as templates, depending on the user’s goals. MSstatsResponse then simulates future experimental observations by adding instances of random variation to the systematic profiles as described in (**Supplementary Algorithm 1**). Each simulated dataset can then be analyzed for (1) target identification (strong vs. weak vs. non-interacting proteins), or (2) OC50 estimation. For OC50 estimation, MSstatsResponse provides the option to estimate OC50 values for each simulated protein and, optionally, to compute confidence intervals for these estimates.

To ensure accurate results, each combination of replicates and doses is repeated in 1,000 simulated datasets. For each combination, MSstatsResponse reports key performance metrics, including average estimated true positive rates for strong and weak interactions and average false positive rates for non-interacting proteins. For OC50 estimation it reports parameter estimates, and option confidence intervals for all simulated proteins. The results are summarized in plots such as in **Fig. 2**, visually trading off doses and replicates.

### Evaluation

#### Evaluation strategy

We compared the performance of MSstatsResponse to that of dr4pl, CurveCurator, and ANOVA. We omitted drc as this older method is relatively similar to dr4pl but less sensitive in drug target identification (**Supplementary Table 2**). Analyses were performed on both experimental and simulated datasets to assess method performance under confirmatory and exploratory experimental designs (**Fig. 1**). Confirmatory experiments were represented by the full benchmark datasets, each with nine doses and three replicates. Exploratory single-replicate experiments were represented by separately analyzed individual replicates from these datasets. Additional analyses generated intermediate designs with varying numbers of doses and replicates.

Each method was evaluated with respect to its ability to address the three analysis tasks in **Fig. 1**. Task 1 (curve fitting) was visually assessed by plotting dose-response curves. Task 2 (target identification) was evaluated with respect to the performance of hypothesis testing. We chose significance thresholds that we believed to be most comparable across methods. Specifically, dr4pl did not provide p-values, but provided parameter estimates and the 95% confidence intervals for all four model parameters: upper asymptote, lower asymptote, slope, and OC50 value. We defined significant drug-protein interactions based on the following criteria: (1) OC50 95% CI is within experimental dose range, and (2) slope is decreasing and 95% CI does not include 0. Evaluations tested using only criteria (1) were overly sensitive and produced variable results across datasets (**Supplementary Fig. 7**). For CurveCurator, we determined drug-target interactions using relevance scores, and users-defined significant thresholds. We defined significance cutoffs as curve fold-change: 0.5 and FDR-adjusted p-value of *q* = 0.05. For ANOVA and MSstatsResponse, evaluations were done at an FDR-adjusted p-value cutoff of *q* = 0.05. Evaluation on the experimental datasets used established Dasatinib targets as positive controls. Evaluation on the simulated datasets was reported in terms of true positive rates 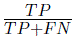 for simulated strong and weak interactors and false positive rates 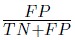 for non-interactors.

Task 3 (OC50 estimation) was evaluated based on the precision of the estimated OC50 values. Precision was measured by the width of their 95% confidence intervals. Consistency of the estimates across different experimental designs was also used to evaluate reproducibility.

### All acquisition strategies were suitable for chemoproteomics experiments

All three acquisition strategies in this study, DIA, TMT, and SRM, produced high-quality data suitable for chemoproteomics experiments. All five known Dasatinib targets demonstrated the expected response patterns (**Supplementary Fig. 8**). While proteome coverage differed across methods (**Table 1**, **Supplementary Fig. 2**), these differences did not substantially limit the detection of drug-protein interactions. The choice of acquisition strategy is most important when considering protein coverage and alignment with the experimental goal. Our benchmark results further indicated that study design and statistical model choice had a greater impact on sensitivity and reproducibility than the acquisition method itself. Specifically, the number of replicates, the selection of drug doses, and the robustness of the statistical model had a much larger influence on performance. These considerations are evaluated in detail in the following sections. We use the DIA benchmark dataset as the primary example in the following sections. Results for TMT and SRM benchmarks are provided in the **Supplementary Sec. 4**.

### In experimental datasets, statistical methods produced distinct dose-response curves

Curve-fit visualizations differed across statistical approaches. dr4pl and CurveCurator both fit four-parameter logistic models and therefore produced nearly identical sigmoidal curves. MSstatsResponse only specified a monotonic non-increasing constraint, which made it robust to non-sigmoidal shapes and resistant to outlier-driven distortions. In contrast, ANOVA modeled each dose independently, which often yielded jagged fits and overfitting to outliers.

**Fig. 3** illustrates these behaviors. Row 1 (SRC, positive control) shows a clean sigmoidal decrease, where all methods agree. Row 2 (CD3G) is flat with a high-dose uptick; dr4pl and CurveCurator forced an increasing sigmoidal shape, ANOVA exaggerated the outlier, and MSstatsResponse maintained a flat, non-increasing fit. Row 3 (YIPF5) is noisy at low doses but decreases at higher doses; all methods except ANOVA captured the overall trend without spurious inflections. Row 4 (SPHK1) rises slightly before decreasing; MSstatsResponse remained flat through the rise and transitioned downward only once the data supported a decline.

**Figure 3:**
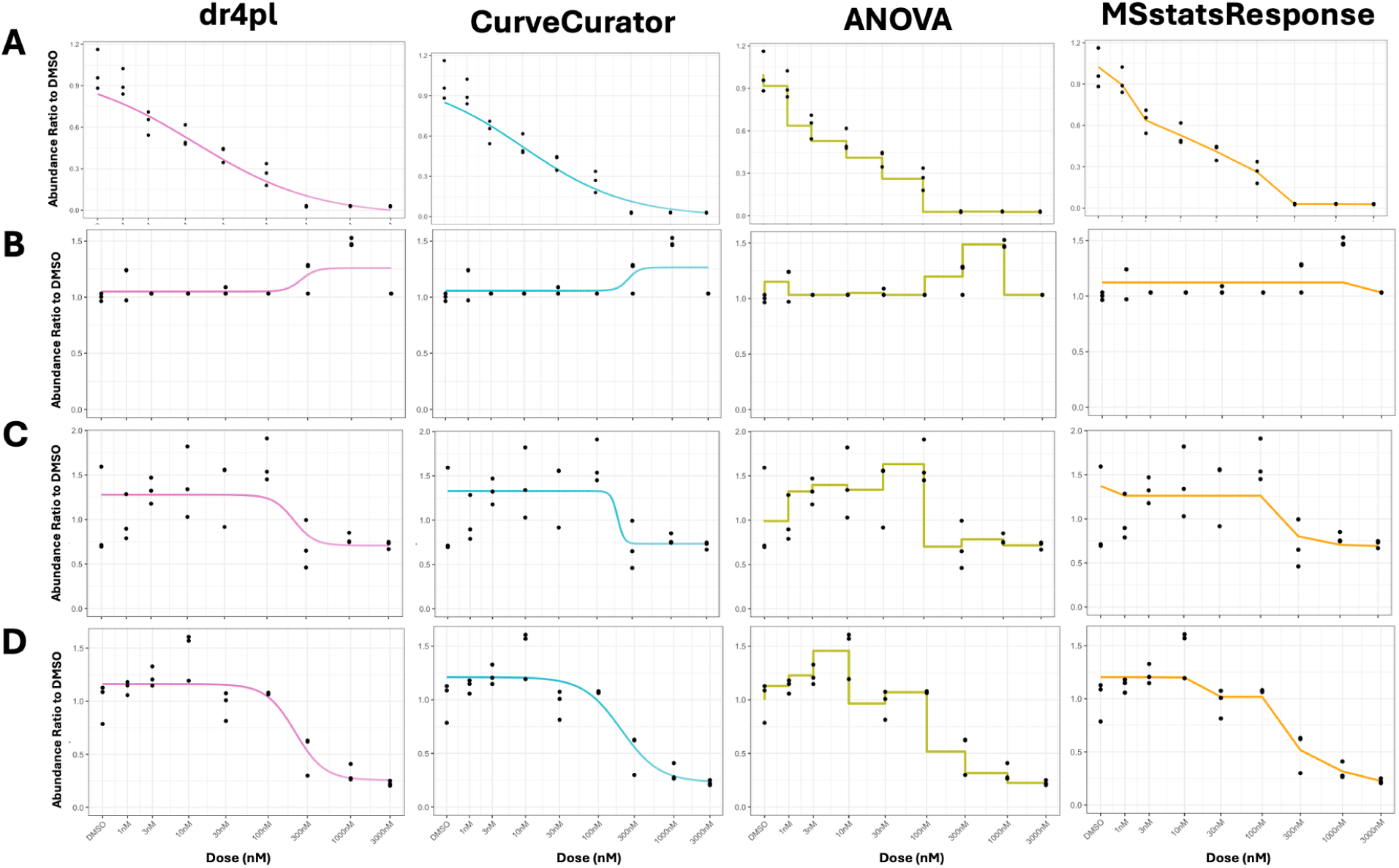
DIA benchmark: examples of fitted dose–response curves across methods. Rows: proteins. Columns: methods/ Dots: input protein abundances. Lines: fitted curves. **(A)** A clear sigmoidal decrease where all methods agree. **(B)** A flat response with a high-dose uptick. dr4pl and CurveCurator impose an increasing sigmoidal shape, ANOVA overfits the outlier, and MSstatsResponse remains flat. **(C)** A noisy profile at low doses with a decrease at higher doses. All methods capture the overall decline except ANOVA, which overfits to the increase. **(D)** Increase at low dose, then decrease at higher doses. MSstatsResponse remains flat through the rise and transitions downward only once the data support a decline.

Similar patterns appeared across other benchmark datasets. In the SRM benchmark (**Supplementary Fig. 8-11**), YES, LCK, and CSK show low-dose increases, where MSstatsResponse remains flat until the decline begins. For clearly sigmoidal profiles (e.g., CSK in DIA and TMT), MSstatsResponse closely matched 4PL fits, recovering sigmoidal shapes when present but not imposing them otherwise. Additional examples for all positive controls are provided in **Supplementary Fig. 8-11**.

### In experimental datasets, statistical methods differed in drug-target detection sensitivity

**Table 3** summarizes the number of significant proteins detected by each method across acquisitions. Across the different acquisition types, the number of detected interactions varied notably by method. No single method detected every positive control in every benchmark dataset. CurveCurator and MSstatsResponse were the closest, while dr4pl failed to detect all the positive controls except for in the DIA dataset. Parametric methods, dr4pl and CurveCurator, missed ABL1 in the SRM dataset (**Supplementary Fig. 9-10**), and MSstatsResponse missed YES in the TMT benchmark with an adjusted p-value of q = 0.066, but detected it in the TMT-RTS suggesting it was borderline (**Supplementary Fig. 8**). CurveCurator detected the most proteins across acquisitions, mostly agreeing with MSstatsResponse, but missed protein GAK exhibiting a weak interaction and picked up some noisier proteins like TNK2 and TGM4 (**Fig. 4**).

**Figure 4:**
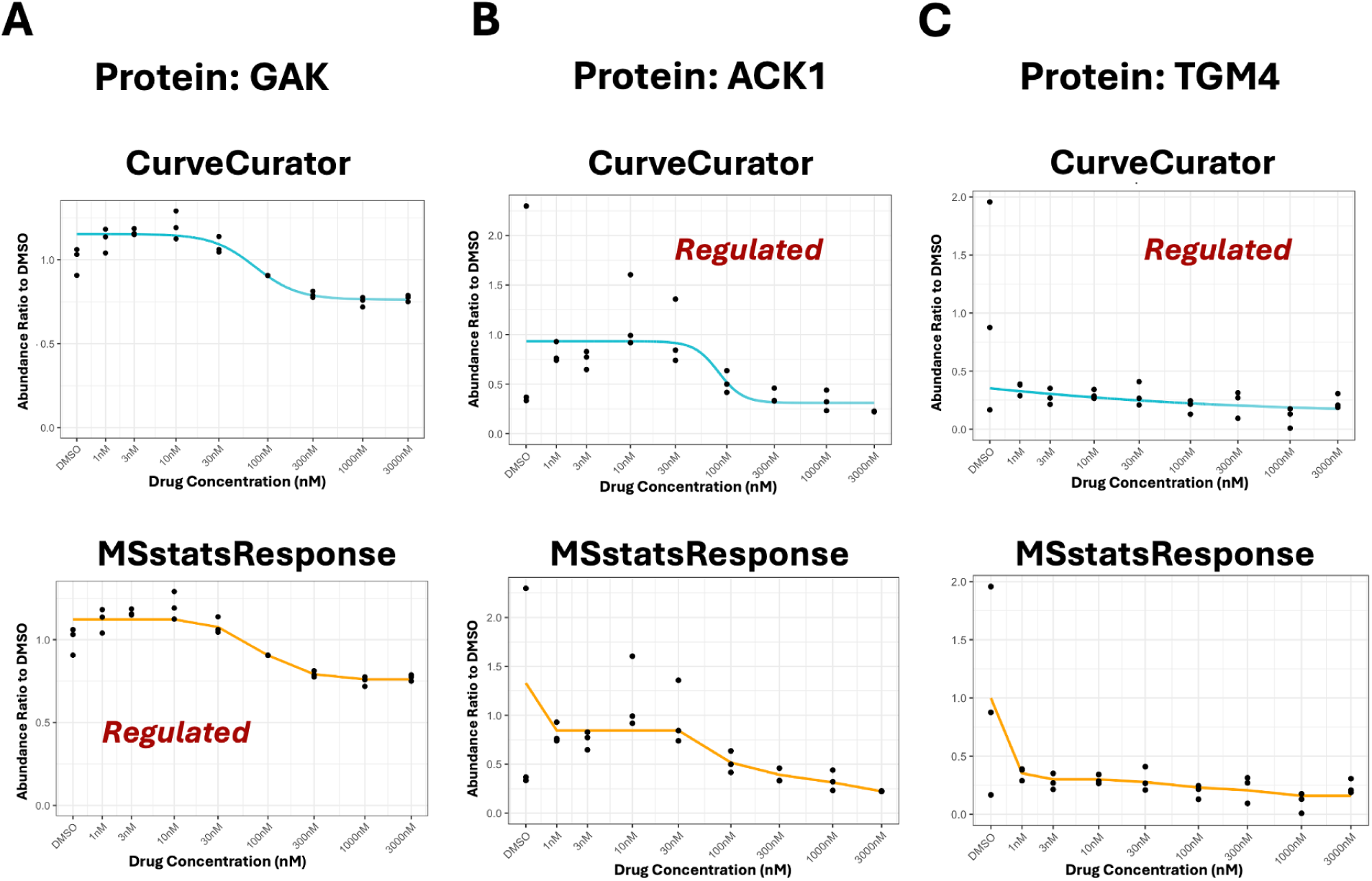
DIA benchmark: comparison of target identification sensitivity across methods. **(A)** MSstatsResponse showed the sensitivity to detect potentially interesting weak/incomplete responses without overfitting to noise. **(B-C)** In contrast, sigmoidal methods such as CurveCurator were more sensitive and occasionally detected noisier profiles like ACK1 and TGM4.

**Table 3:** Benchmark dataset significant protein-drug interactions across acquisition types and methods using all replicates. Results using full datasets in DIA, TMT, TMT with RTS, and SRM datasets. The number of significant proteins is shown for each method. Known Dasatinib targets (SRC, ABL1, YES, CSK, and LCK) are shown in parentheses and serve as positive controls.

|  | **DIA** | **SRM** | **TMT** | **TMT-RTS** |
| --- | --- | --- | --- | --- |
| dr4pl | 9 (5) | 4 (4) | 6 (3) | 7 (3) |
| CurveCurator | 31 (5) | 6 (4) | 13 (5) | 11 (5) |
| ANOVA | 8 (5) | 5 (5) | 6 (4) | 10 (4) |
| MSstatsResponse | 14 (5) | 5 (5) | 9 (4) | 11 (5) |

### In pseudo-single-replicate experimental datasets, drug-target detection varied between replicates

Single-replicate experimental designs present a challenge for detecting drug-protein interactions using dose-response models. Parametric curve fitting methods are particularly sensitive to variability in the input data. Without replicates, small fluctuations or outliers at a single dose can substantially change the estimated curve shape, resulting in misleading dose-response representations and missed interactions.

To demonstrate this in practice, we analyzed each replicate from our triple-replicate DIA benchmark experiment independently to mimic three “pseudo” single-replicate design experiments. **Fig. 5** shows that the results varied greatly depending on which replicate was used, demonstrating the lack of reproducibility in single-replicate experiments. dr4pl and drc failed to pick up all positive controls. Although CurveCurator picked up all the positive controls, the agreement between replicates analyzed varied substantially, with as many as 66 unique proteins detected in the analysis of replicate 2. These patterns were further exaggerated in the other benchmark experiments, where all parametric methods (i.e., dr4pl, drc, CurveCurator) failed to detect every positive control in each replicate (**Supplementary Fig. 12-14**).

**Figure 5:**
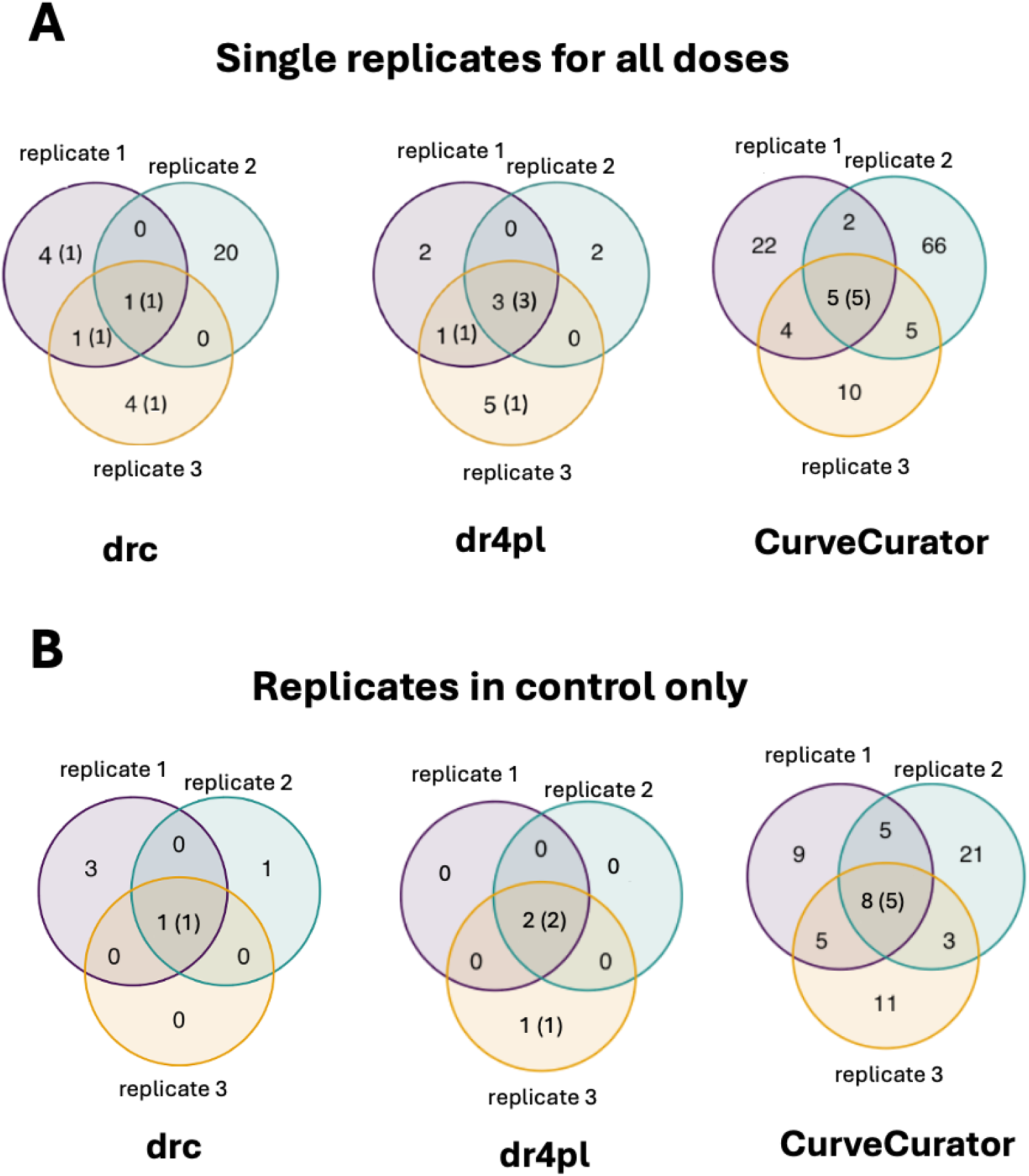
DIA benchmark: replicate choice influenced detection of drug-protein interactions. **(A)** Single replicate per dose. Rows show drc, dr4pl, and CurveCurator (top to bottom). Analyses using only one replicate at each dose produced unstable detection: drc showed low sensitivity (CSK most consistently detected), dr4pl correctly identified three known Dasatinib targets (SRC, ABL1, CSK) while only replicate 3 recovered all five targets (SRC, ABL1, CSK, YES, LCK). CurveCurator recovered all five targets but also detected many additional proteins with substantial between-replicate variability. **(B)** Replicates in DMSO only. Inclusion of replicates in the control reduced random detections for all methods relative to (A). Overall sensitivity remained limited.

To demonstrate the benefit of including replicates in the control condition, we repeated the analysis using all three DMSO replicates. As shown in **Fig. 5B**, adding control replicates reduced the number of detected proteins significantly for all replicate groups, indicating that the inclusion of replicates in the DMSO condition can help reduce noise and improve reproducibility. However, overall sensitivity remained limited in methods such as drc and dr4pl without replicates in all doses.

### In simulated datasets, MSstatsResponse detected weak interactions without increasing false positives

All statistical methods effectively detect strong drug-protein interactions under ideal experimental conditions (i.e., with low variability, a high number of doses, and multiple replicates per dose). However, semi-parametric approaches, MSstatsResponse and ANOVA, show promise for increased sensitivity in identifying weak interactions while maintaining control over false positives.

**Fig. 6** demonstrates this in practice by evaluating each method’s performance on simulated strong, weak, and non-interacting protein profiles with a triple-replicate, 9-dose experimental design derived from the experimental DIA dataset. Semi-parametric methods, MSstatsResponse and ANOVA, had high recall and sensitivity by picking up all strong interactions, and majority of weak interacting proteins, while MSstatsResponse did so without increasing the false positive rate for non-interacting proteins. This showed that MSstatsResponse has increased sensitivity, while controlling the false-positive rate.

**Figure 6:**
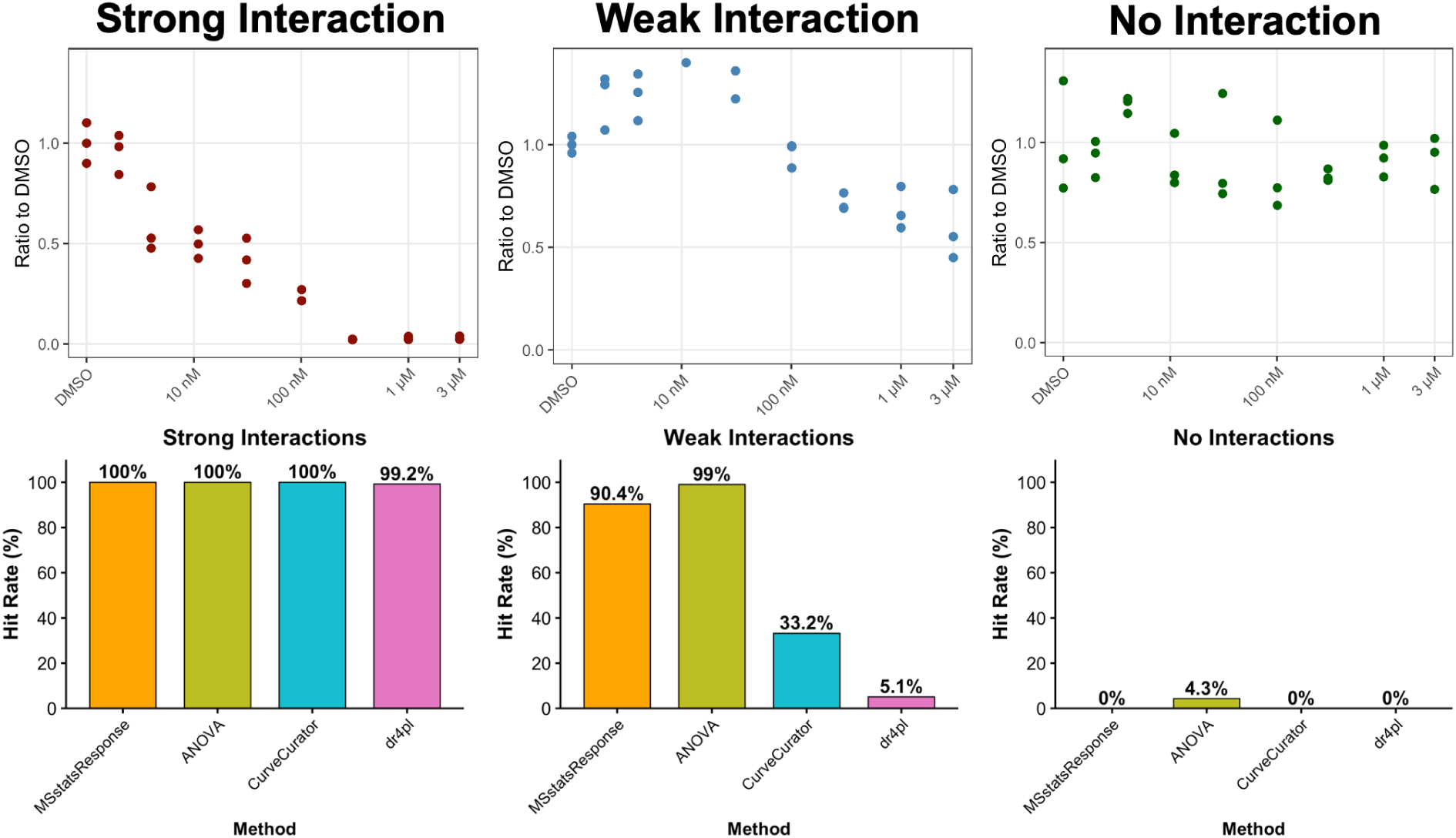
Simulated dataset: semi-parametric methods, ANOVA and MSstatsResponse, achieved best target identification detection of weakly interacting proteins. **(A-C)** Simulated protein response profile examples based on real DIA-derived templates. **(A)** Strong interacting protein simulated using the SRC response profile. **(B)** Weak interacting protein simulated using the TEC response profile. **(C)** Non-interacting protein simulated using EIF2AK4 profile. Each bar chart shows the hit rate across 1,000 simulation runs for a given profile, based on 9 doses (DMSO, 1 nM, 3 nM, 10 nM, 30 nM, 100 nM, 300 nM, 1000 nM, 3000 nM), 3 replicates per dose, and a variance of 0.4. All methods perform well under ideal conditions. MSstatsResponse and ANOVA demonstrate high sensitivity in detecting both strong and weak interactions, while MSstatsResponse does so without compromising specificity (i.e., no false positives).

When data variability increased, all methods declined in sensitivity for detecting weak interactions (**Supplementary Fig. 15**). Notably, the performance of dr4pl drops substantially under noisy conditions, and failed to detect almost all strong interactions. This highlights the robustness of semi-parametric methods, and the dependence of parametric methods on high data quality.

### In simulated datasets with fewer doses, MSstatsResponse maintained drug-target detection sensitivity

Experimental designs with less doses are sometimes necessary in chemoproteomics due to cost and sample space limitations. To evaluate method performance under such constraints, we simulated a dataset with four dose conditions (DMSO, 1 nM, 1000 nM, 3000 nM), each with three replicates, across 2,000 proteins with a variance of 0.4.

**Fig. 7** shows that MSstatsResponse and ANOVA maintained high sensitivity for strong interactions despite reduced dosage coverage. CurveCurator showed moderate sensitivity but began to lose power under these conditions, while dr4pl failed to detect strong interactions entirely. These results demonstrate that classic parametric curve fitting methods require many doses to perform effectively. MSstatsResponse, by contrast, effectively controlled false positives through its monotonicity constraint, whereas ANOVA was more sensitive to outliers, leading to inflated false-positive rates among non-interacting proteins.

**Figure 7:**
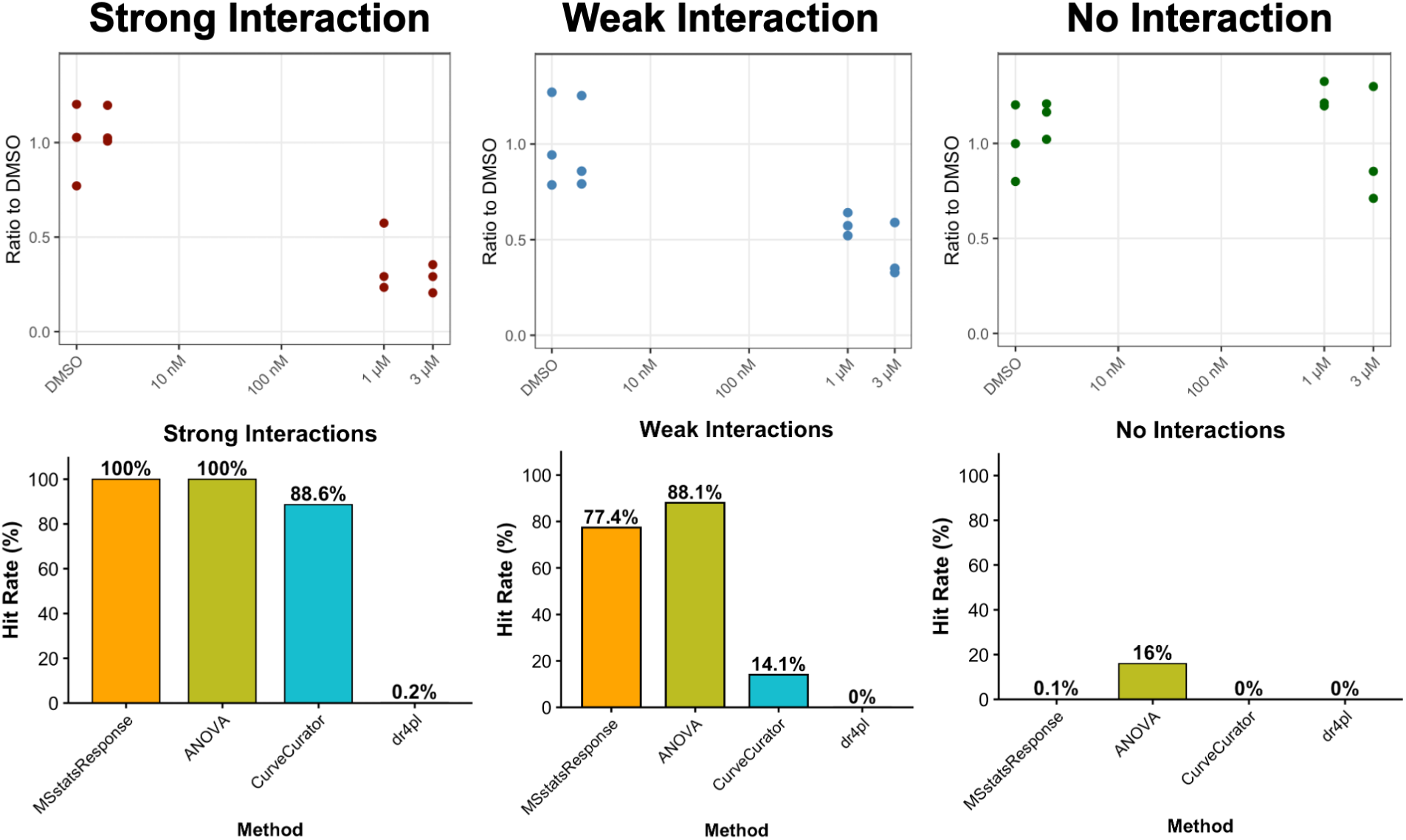
Simulated dataset: MSstatsResponse maintained high sensitivity for strong protein interaction detection without increased false positive rate for datasets with fewer doses. **(A)** Method performance evaluated on a simulated dataset with 4 doses (DMSO, 1 nM, 1000 nM, 3000 nM), 3 replicates per dose, and a variance of 0.4 across 2,000 proteins. MSstatsResponse and ANOVA maintain high sensitivity despite fewer dose points. CurveCurator retains moderate sensitivity but begins to lose power under reduced dose designs. dr4pl is unable to operate on datasets with only four dose levels due to model fitting constraints. **(B)** Specificity analysis for non-interacting proteins showed that ANOVA is sensitive to dose condition outliers, leading to false positives. In contrast, MSstatsResponse retains both high sensitivity and specificity by the model’s monotonicity constraint.

To generalize beyond a single simulated design, we generated power curves across a range of scenarios (**Fig. 8**). These curves show that strong interactions can be detected with relatively few doses, provided replicates are included, while weak interactions require both increased dose levels and replication for reliable detection. To assess whether these trends extend to experimental data, we applied the same analysis framework to benchmark datasets with fewer dose levels. In this setting, target detection patterns were broadly comparable across acquisition types (**Supplementary Table 3**). For experiments involving a single high-dose treatment, we recommend standard differential abundance methods (e.g., MSstats^40^), which are better suited for single-dose target identification. Overall, these results demonstrate that semi-parametric approaches like MSstatsResponse can maintain robust performance with fewer doses, offering a practical alternative design strategy when resources are limited.

**Figure 8:**
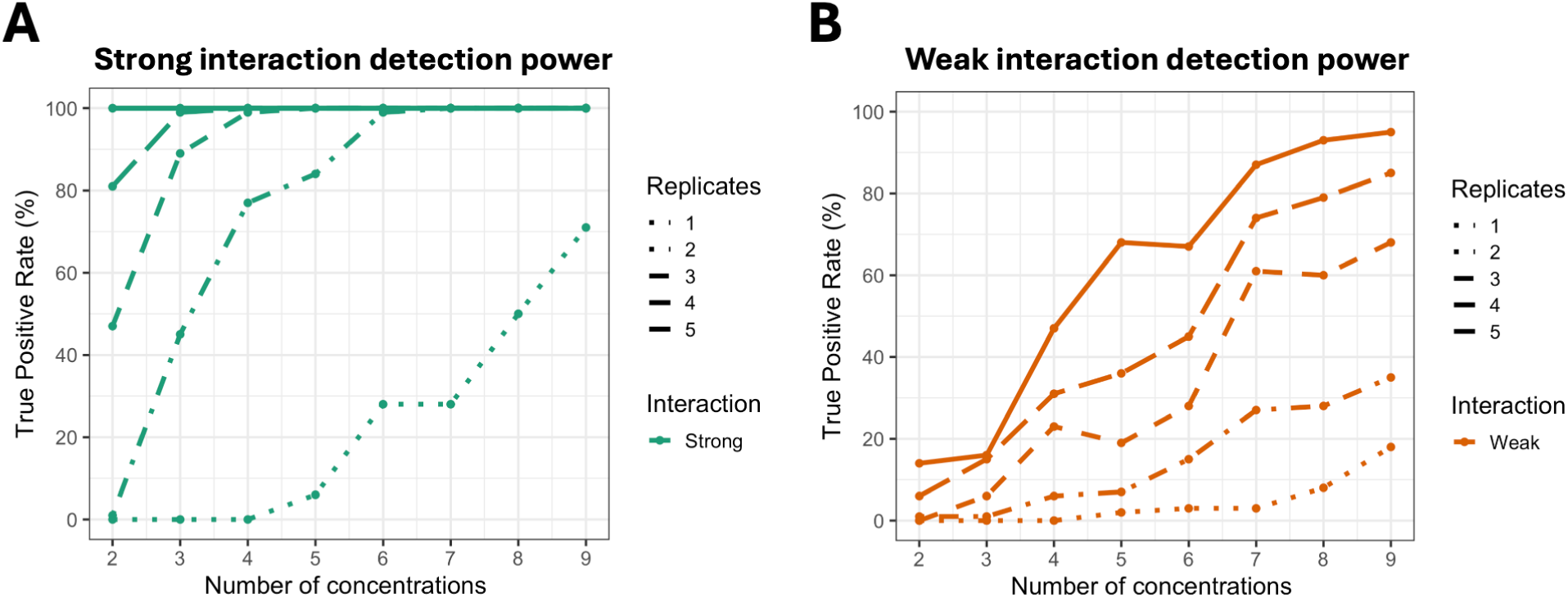
Simulated Dataset: detected interactions depended on the number of doses and replicates per dose. Simulated dose–response experiments were generated using increasing numbers of concentrations (in nM): 2-dose (0, 3000), 3-dose (0, 1000, 3000), 4-dose (0, 1, 1000, 3000), 5-dose (0, 1, 100, 1000, 3000), 6-dose (0, 1, 100, 300, 1000, 3000), 7-dose (0, 1, 10, 100, 300, 1000, 3000), 8-dose (0, 1, 10, 30, 100, 300, 1000, 3000), and 9-dose (0, 1, 3, 10, 30, 100, 300, 1000, 3000). **(A)** Detection of strong interactions (i.e., proteins whose probe-bound abundance decreases to near zero at the highest dose) approached 100% with 5 replicates and as few as 2 doses. Experiments with 6+ doses required at least two replicates for reliable detection. **(B)** Detection of weak interactions (i.e., proteins whose probe-bound abundance decreases to roughly 50% of the control level) required more replicates and more doses.

### In experimental datasets, MSstatsResponse produced precise OC50 estimates

Among the methods compared, MSstatsResponse and dr4pl provide confidence intervals for OC50 estimation, whereas CurveCurator produces only a single point estimate and ANOVA does not support this parameter at all. We therefore focused our evaluation on MSstatsResponse and dr4pl, and included CurveCurator’s point estimates for reference.

We compared these methods across three experimental design scenarios (all with three replicates per dose): a full 9-dose design (DMSO + 8 doses), a reduced 5-dose design (2 low, 1 middle, 2 high), and minimal 4-dose design (2 low, 2 high).

**Fig. 9** shows OC50 estimates and 95% confidence intervals for the drug target ABL1 under each design. While both methods yielded similar results with the full-dose experiment, dr4pl produced increasingly wide and unstable confidence intervals as doses were removed. This instability stems from dr4pl’s reliance on a sigmoidal dose-response assumption, which cannot be reliably fit with limited dose points. CurveCurator generated a single OC50 estimate in each scenario, but without confidence intervals its reproducibility and statistical interpretability were limited.

**Figure 9:**
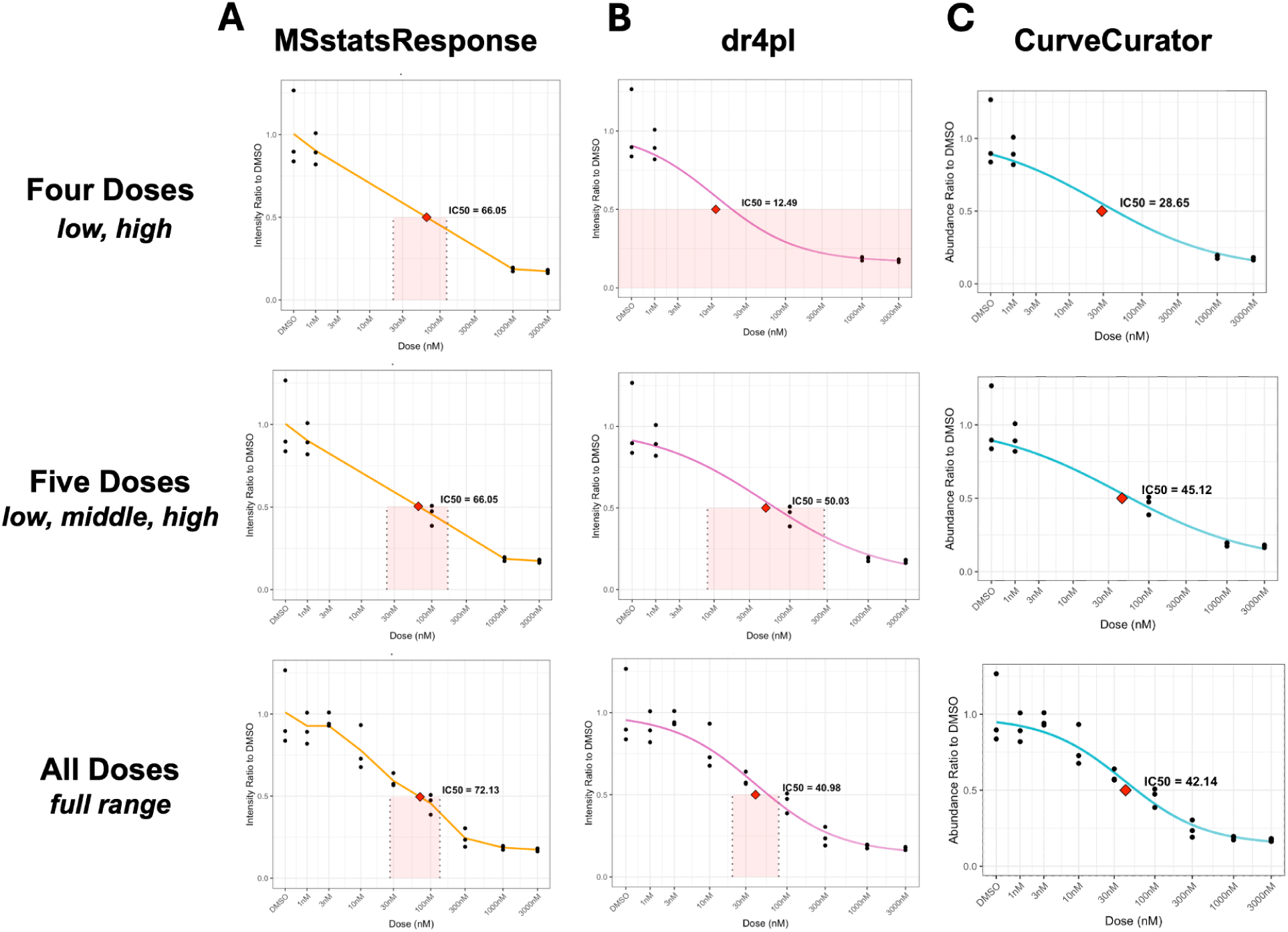
DIA benchmark: MSstatsResponse maintained stable OC50 estimates across experimental designs with fewer doses. Columns compare three estimation methods, MSstatsResponse, dr4pl, and CurveCurator, across varying experimental designs. **(A)** MSstatsResponse produced consistent OC50 estimates with narrow bootstrapped confidence intervals across four-, five-, and nine-dose designs. **(B)** dr4pl required more dose points to achieve stability. Without a middle dose, confidence intervals were wide or unrealistic. **(C)** CurveCurator generated stable single OC50 point estimates in all designs. No confidence intervals are provided to assess uncertainty.

In contrast, MSstatsResponse combined isotonic regression with linear interpolation and bootstrapping to generate stable and reproducible OC50 estimates with valid confidence intervals under all scenarios. Because it assumes only that dose–response relationships are monotonic (non-increasing), rather than strictly sigmoidal, MSstatsResponse remained robust even when the number of available doses was limited. These results highlight its advantage in experimental settings where cost or sample constraints restrict dose coverage. Despite these results, we still recommend including as many dose levels as feasible when the experimental goal is precise and reliable OC50 estimation.

## Discussion

We propose MSstatsResponse, a semi-parametric statistical framework for the analysis of chemoproteomics experiments. Our results demonstrate that MSstatsResponse is well positioned to analyze large-scale proteomics drug screens across a range of experimental designs and acquisition strategies. Unlike fully parametric models, which fit curves of pre-specified shape and required many doses and replicates to achieve stability, MSstatsResponse was able to maintain sensitivity in more resource-limited settings.

A key finding of this study is that experimental design choice strongly influenced the reliability of dose–response analysis results. While large-scale screens often favor single-replicate, multi-dose designs, we showed that these designs were highly sensitive to experimental noise and exhibited poor reproducibility (**Fig. 5**). In these cases, parametric models frequently overfit or misinterpreted random fluctuations which lead to increased false positives. In contrast, a minimal dose, multi-replicate design provided a more reliable foundation for initial hit detection. We therefore recommend this design as a starting point for exploratory screens, followed by targeted dose–response profiling to assess potency.

We recommend at the very least, for exploratory screens to include replicates in the control condition. Without replicated controls, ratio-scale normalization can amplify unseen variability in the control condition which can artificially distort downstream dose–response measurements. We show in **Fig. 5** that the inclusion of replicates in the control condition can at the very least control the false-positive rate and limit sensitivity to random fluctuations in the dataset.

For confirmatory analyses, adding multiple doses with replication provided the strongest foundation for both interaction detection and OC50 estimation. As demonstrated in our simulations and benchmarks, this design enabled more accurate curve fitting, increased sensitivity to weak interactions, and more stable parameter estimates (**Fig. 6**, **Fig. 9**). Although this option is more resource-intensive, this design is strongly recommended when the analysis goal is high-confidence detection and potency estimation.

We recognize that reducing the number of drug doses is sometimes necessary due to cost limitations. We found that MSstatsResponse retained robust performance with as few as four doses with replicates (**Fig. 7**). In contrast, fully parametric models such as dr4pl often failed under reduced designs, as the limited dose range was insufficient to fit all curve parameters reliably.

We further showed that all acquisition strategies tested (i.e., DIA, DDA, and SRM) were suitable for chemoproteomics dose–response analysis (**Table 3**). We conclude that data quality should not be a concern and acquisition method selection can be guided primarily by instrument availability and throughput requirements.

Altogether, our findings lead to three major recommendations. First, when resources are limited, we recommend prioritizing biological replicates while limiting the number of doses. Replicates consistently improved robustness across all methods and tasks and were essential for reliable inference. Second, when possible, combine replication with at least one intermediate and one high dose to enable flexible curve modeling and accurate OC50 estimation. MSstatsResponse is particularly well suited to these designs, as it provided robust inference without relying on strict curve assumptions. And third, if deciding to do a single-replicate dose response screen, we strongly recommend the inclusion of replicates in the control condition to stabilize baseline estimates, reduce false-positives, and further increase screen reproducibility.

Beyond chemoproteomics, the MSstatsResponse framework is readily extendable to a wide range of dose-response–like studies. These include transcriptomics dose-response experiments, thermal profiling experiments where temperature replaces dose, protein turnover studies, or more generally, any biological relationship that follows a monotonic trend. In this study we focused on non-increasing constraints, reflecting the behavior of inhibitors, but MSstatsResponse can be applied equally well to datasets where responses are expected to be non-decreasing. The underlying methods and workflow remain unchanged. Future work will focus on the integration of MSstatsResponse into the MSstatsShiny online GUI which will streamline data processing, analysis and data visualization for users.

## Data and code availability

The four benchmark datasets analyzed in this manuscript are publicly available on MassIVE under the accession numbers MSV000100970 (DIA dose-response dataset), MSV000100968 (TMT dose-response datasets), and MSV000100969 (SRM dose-response dataset). The intermediate datasets and analysis scripts used to reproduce the results in this manuscript are available on Zenodo at https://doi.org/10.5281/zenodo.18881723. The MSstatsResponse R package is available through Bioconductor at https://bioconductor.org/ packages/MSstatsResponse.

