## Supplemental Materials for "MSstatsResponse: Semi-parametric statistical model enhances detection of drug-protein interactions in chemoproteomics experiments"

#### Supplementary Information

Sarah Szvetecz[1,2] Devon Kohler[1,2] S. Denise Field[4] Pierre Jean-Beltran[4]
Hyunsuk Suh[3] Robert J. Seward[3] Joel D. Federspiel[3] Liang Xue[4]
Olga Vitek[1,2*]

[1]Khoury College of Computer Sciences, Northeastern University, Boston, MA, USA

[2]Barnett Institute for Chemical and Biological Analysis, Northeastern University, Boston, MA, USA 02115

[3]Pfizer Inc., Andover, MA, USA

[4]Pfizer Inc., Cambridge, MA, USA

### 1 Experimental datasets

#### 1.1 Benchmark experiments

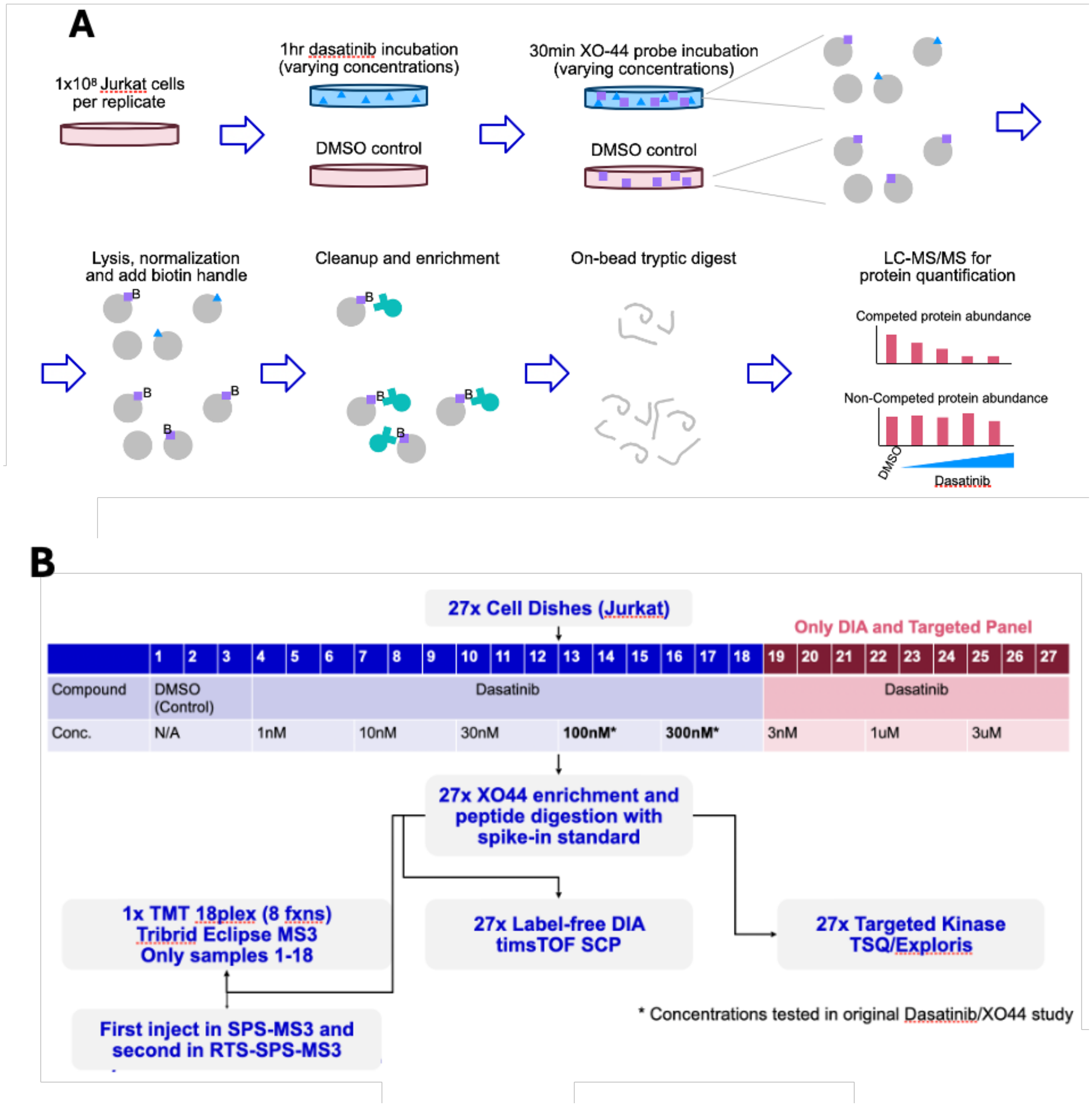

Figure 1: **Benchmark datasets: experimental design.** **(A)** Cells were treated with varying doses of Dasatinib in competition with the XO44 pan-kinase probe and analyzed using data-independent acquisition (label-free DIA), data-dependent acquisition (TMT 18-plex, standard MS3 and real-time search), and targeted selected reaction monitoring (SRM). **(B)** All datasets included triplicate biological replicates per condition. TMT datasets included five drug doses (1 nM, 10 nM, 30 nM, 100 nM, and 300 nM) plus DMSO controls. DIA and SRM datasets included eight drug doses (1 nM, 3 nM, 10 nM, 30 nM, 100 nM, 300 nM, 1 $\mu$M, and 3 $\mu$M) plus DMSO controls.

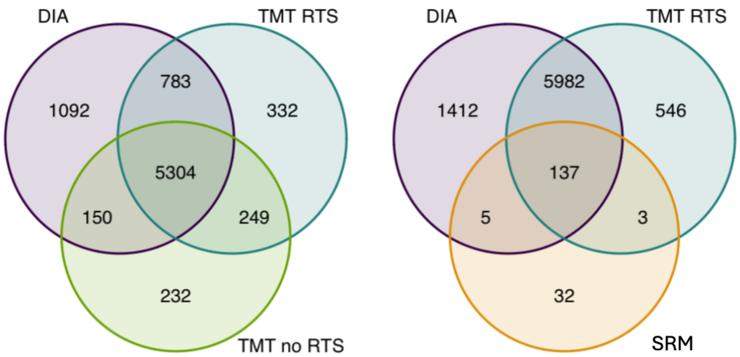

Figure 2: **Benchmark datasets: number of proteins identied per acquisition.** All datasets successfully identified the five known Dasatinib targets (SRC, ABL1, YES, LCK, and CSK) and the negative control EIF2AK4. Real-time search improved peptide identification in the TMT workflow, increasing protein coverage by 733 proteins relative to standard MS3 acquisition.

#### 1.2 Simulated datasets

The simulations of log-scale protein abundances used as a template the abundances of a selected protein from a past dose-response experiment. In this manuscript, the template proteins were chosen from the DIA benchmark dataset to represent strong interactions, weak interactions or no interactions. The corresponding template abundances were derived by averaging the log2-transformed protein intensities across replicates at each dose level.

Additional variation was added at each dose level, to introduce randomness and create the number of desired replicates. In this manuscript, the variation was estimated from the average protein-level variance across sample groups in the DIA benchmark dataset.

Outliers were optionally added to some simulations by adding values from the log-normal distribution. In this manuscript, the probability of an outlier was set to 0 as default, and set to 5% when investigating outliers.

**Algorithm 1** Simulation of dose–response Data

**Input:**
$D$: number of drug doses (excluding control)
$R$: number of replicates per dose
$\sigma^2$: technical and biological variance
$\{\mu_d, d = 0, \ldots, D\}$: template protein abundances (e.g., strong, weak, no interaction) per dose
$p_{\text{outlier}}$: probability of an outlier

**Output:**
$\{y_{dr}, d = 0, \ldots, D;\ r = 1, \ldots, R\}$: simulated protein-level log-abundances across doses and replicates

**for** $d = 0$ to $D$ **do**
  **for** $r = 1$ to $R$ **do**
    $y_{dr} \leftarrow \mu_d + \mathcal{N}(0, \sigma^2)$ ▷ Add noise to template value

    $isMissing \leftarrow Bernoulli(p_{\text{outlier}})$ ▷ Random outlier selection
    **if** $isMissing = 1$ **then**
      $s \leftarrow 2 \cdot Bernoulli(0.5) - 1$ ▷ Random direction selection
      $y_{dr} \leftarrow y_{dr} + s \cdot \text{LogNormal}(0, 1)$ ▷ Apply log-Normal deviation
    **end if**
  **end for**
**end for**
**Output:** $\{y_{dr}, d = 0, \ldots, D;\ r = 1, \ldots, R\}$

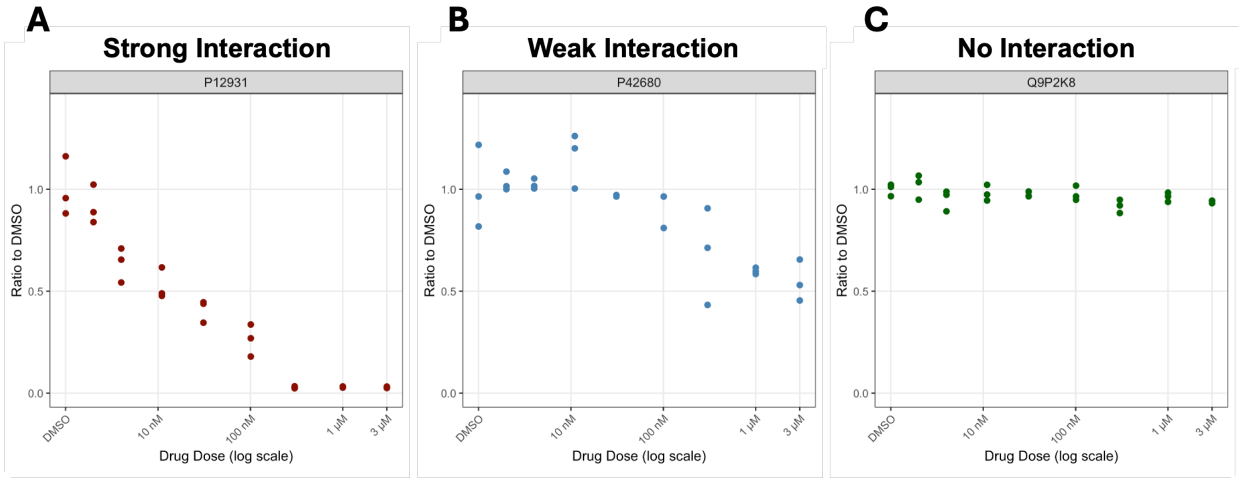

Figure 3: **DIA benchmark: protein templates chosen for simulation.** **(A)** SRC protein, strong interaction. Strong interactions served as templates for full inhibition, where protein abundance decreased to near zero at the highest drug dose. **(B)** TEC protein, weak interaction. Weak interactions represented partial inhibition, where protein abundance decreased to approximately 50% of baseline levels. **(C)** EIF2AK4 protein, no interaction. Non-interacting proteins maintained constant abundance across all doses, indicating no drug effect.

### 2 Background

| Model | Hypothesis testing | Reject $H_0$ | Degrees of freedom | OC50 |
|---|---|---|---|---|
| **drc** | $H_0: \theta_3 = 0 \quad \text{or} \quad \theta_2 \notin [x_{\min}, x_{\max}]$<br>$H_a: y_{dr} = \theta_1 + \frac{\theta_4 - \theta_1}{1 + 10^{\theta_3(x_d - \log_{10}\theta_2)}} + \epsilon_{dr}$<br>$\epsilon_{dr} \overset{\text{iid}}{\sim} N(0, \sigma^2)$ | $\hat{\theta}_3 < 0$ and<br>$\hat{\theta_2} \in [x_{\min}, x_{\max}]$ and<br>p-value for $\theta_3$ and $\theta_2 < \alpha$ | N/A | $\theta_2$ |
| **dr4pl** | $H_0: \theta_3 = 0 \quad \text{or} \quad \theta_2 \notin [x_{\min}, x_{\max}]$<br>$H_a: y_{dr} = \theta_1 + \frac{\theta_4 - \theta_1}{1 + 10^{\theta_3(x_d - \log_{10}\theta_2)}} + \epsilon_{dr}$<br>$\epsilon_{dr} \overset{\text{iid}}{\sim} N(0, \sigma^2)$ | $\hat{\theta}_3 < 0$ and<br>$\hat{\theta}_2 \in [x_{\min}, x_{\max}]$ and<br>95% CI for $\theta_2$ excludes 0 | N/A | $\theta_2$ |
| **CurveCurator** | $H_0: y_{dr} = 0x + 1 + \epsilon_{dr}$<br>$H_a: y_{dr} = \theta_1 + \frac{\theta_4 - \theta_1}{1 + 10^{\theta_3(x_d - \log_{10}\theta_2)}} + \epsilon_{dr}$<br>$\epsilon_{dr} \overset{\text{iid}}{\sim} N(0, \sigma^2)$ | $F_{obs} = \frac{\text{SSE}_0 - \text{SSE}_a}{\text{SSE}_a} \cdot \frac{DR}{4}$<br>$> F(1-\alpha, df_1, df_2)$<br>and $\lvert \hat{y}'_{x_{\min}} - \hat{y}'_{x_{\max}} \rvert > \delta$ | $df_1 = 5$<br>$df_2 = (0.8 - c) \times$<br>$(DR - 2.5)$, where<br>$c = \frac{1}{\frac{(DR-4)^4}{DR} + 4}$ | $\theta_2$ |
| **ANOVA** | $H_0: y'_{dr} = \mu + \epsilon_{dr}$<br>$H_a: y'_{dr} = \mu + x_d + \epsilon_{dr}$<br>$\sum_{d=0}^{D} x_d = 0, \quad \epsilon_{dr} \overset{\text{iid}}{\sim} N(0, \sigma^2)$ | $F_{obs} = \frac{\frac{\text{SSE}_0 - \text{SSE}_a}{df_0 - df_a}}{\frac{\text{SSE}_a}{df_a}}$<br>$> F(1-\alpha, df_0 - df_a, df_a)$ | $df_0 = (D+1)R - 1$<br>$df_a = (D+1)(R-1)$ | N/A |
| **MSstatsResponse** | $H_0: y'_{dr} = \mu + \epsilon_{dr}$<br>$H_a: y'_{dr} = f(x_d) + \epsilon_{dr}$<br>$\epsilon_{dr} \overset{\text{iid}}{\sim} N(0, \sigma^2)$, where<br>$f(x_0) > f(x_1) > \cdots > f(x_D)$ | $F_{obs} = \frac{\frac{\text{SSE}_0 - \text{SSE}_a}{df_0 - df_a}}{\frac{\text{SSE}_a}{df_a}}$<br>$> F(1-\alpha, df_0 - df_a, df_a)$ | $df_0 = (D+1)R - 1$<br>$df_a = (D+1)R - y^*$ | N/A |
| | $H_0: y_{dr} = \mu + \epsilon_{dr}$<br>$H_a: y_{dr} = f(x_d) + \epsilon_{dr}$<br>$\epsilon_{dr} \overset{\text{iid}}{\sim} N(0, \sigma^2)$, where<br>$f(x_0) > f(x_1) > \cdots > f(x_D)$ | N/A | N/A | $x_d :$<br>$f(x_d) =$<br>0.5 |

Table 1: **Summary of statistical modeling and inference for one protein, used for task 2 (target identification at level $\alpha$), and task 3 (OC50 estimation).** Notation is as in the main text. $y_{dr}$ are protein abundances on the ratioed scale; $y'_{dr}$ are protein abundances on $\log_2$-transformed (or compatible) scale. $D$ is the number of doses excluding the control. The table assumes an equal number of replicates $R$ for each dose. $\text{SSE}_0$ and $\text{SSE}_a$ are the sum of squares of the error of the null and alternative models, respectively. $F$ is the probability distribution. $\hat{}$ refers to values estimated from the data. $y*$ is the number of unique fitted values used to fit the isotonic regression model. In proteins where the monotonicity constraints are not broken, $y^* = D + 1$. For convenience of interpretation, MSstatsResposne fits the isotonic regression on the $\log_2$-transformed scale for task 2, and on the ratioed scale for task 3. However, OC50 can be equivalently estimated from the log-scale fit as $\log_2\left(\frac{1}{2}\hat{y}_{0\cdot})\right) = \hat{f}(x_0) - 1$.

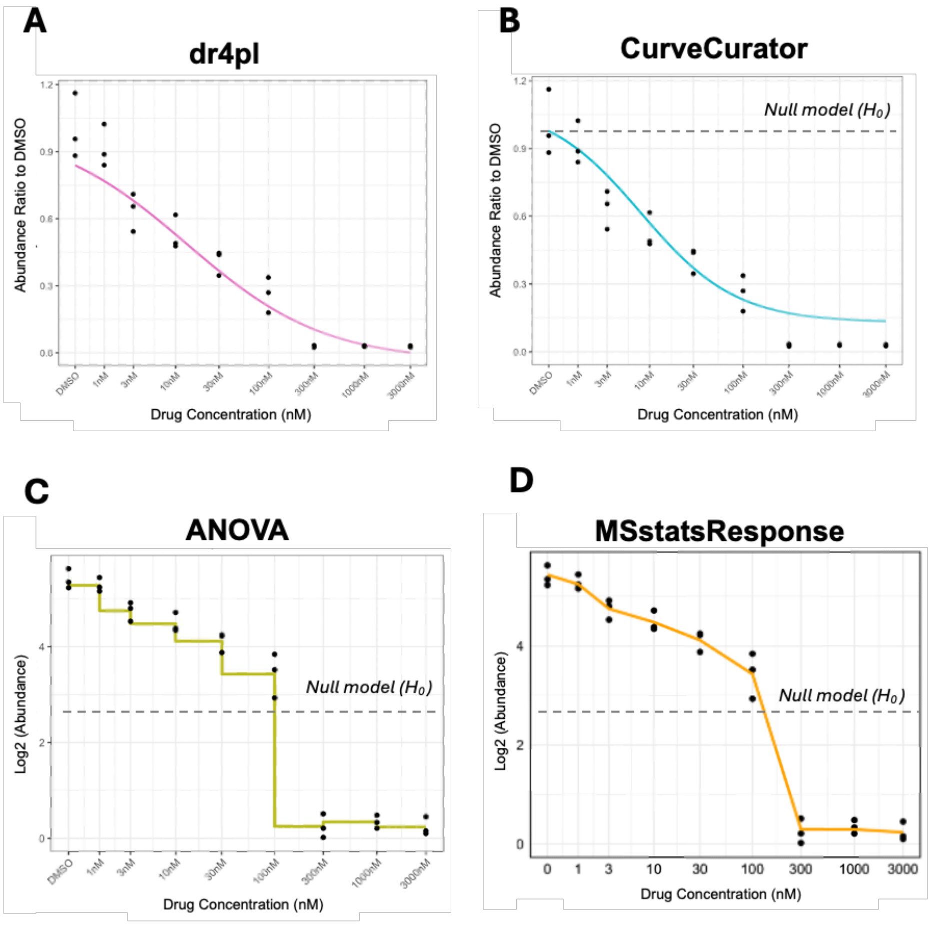

Figure 4: **DIA benchmark, positive control protein SRC.** Dose-response curves for SRC, a known Dasatinib target, are shown for two parametric methods(**(A)** dr4pl, **(B)** CurveCurator) and two semi-parametric (**(C)** ANOVA, **(D)** MSstatsResponse). Each panel displays the fitted model curve and the null hypothesis used for target identification.

### 3 Results

#### 3.1 Overall workflow in MSstatsResponse

##### Task 2: Target identification

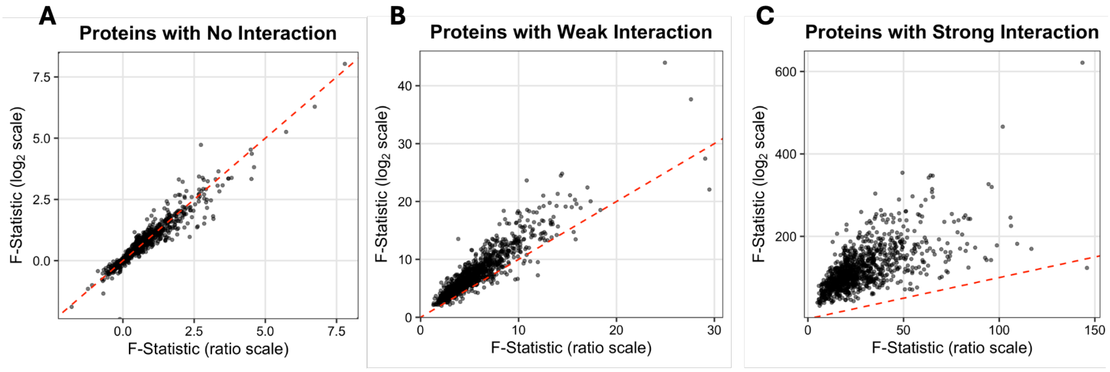

Figure 5: **Simulated dataset: target identification with MSstatsResponse on the log scale increased detection sensitivity without increasing false positives.** Simulation datasets containing 3,000 proteins (n = 1,000 in each strength group) were used to compare detection sensitivity on the log scale versus the ratio scale. **(A)** F-statistics for negative controls were similar on both scales. **(B)** Proteins with weak interactions showed slightly larger F-statistics on the log scale compared to the ratio scale. **(C)** Similar patterns were observed for proteins with strong interactions, although all F-statistics were sufficiently large to identify significance on both scales.

##### Task 3: OC50 estimation

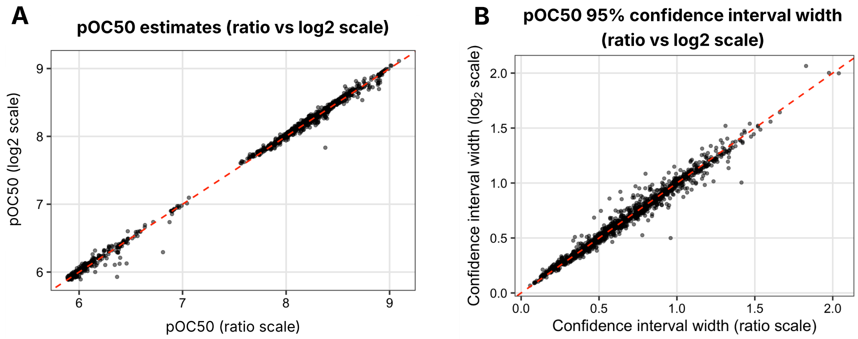

Figure 6: **Simulated dataset: OC50 estimates from MSstatsResponse are near identical on both the log and ratio scale.** OC50 values are reported as pOC50 ($-\log_{10}$OC50), the default output of MSstatsResponse. Simulation datasets containing 3,000 proteins (n = 1,000 in each strength group) were used to compare pOC50 estimates and confidence interval widths on both scales. **(A)** pOC50 estimates follow a 1:1 trend regardless of what scale the data used to fit the model was. **(B)** shows that a similar trend was observed in 95% confidence interval widths.

### 4 Evaluation

#### 4.1 Evaluation strategy

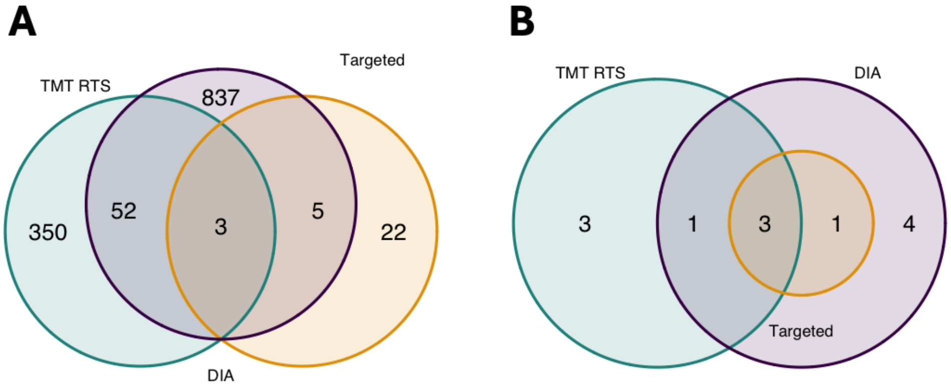

Figure 7: **DIA benchmark: dr4pl results were sensitive to how drug–protein interactions were defined.** (**A**) Using the OC50 estimate alone (i.e., requiring only that the OC50 95% confidence interval fall within the experimental dose range) produced a high number of false positives and substantial variability in the number of hits called across acquisitions. (**B**) A combined criterion that required both an in-range OC50 95% CI and a decreasing slope whose 95% CI excludes 0, produced more stable and reliable detection results. Three positive controls (SRC, CSK, LCK) were consistently detected across all acquisitions, while GAK was shared between TMT and DIA, and YES was shared between Targeted and DIA.

#### 4.2 All acquisition strategies were suitable for chemoproteomics experiments

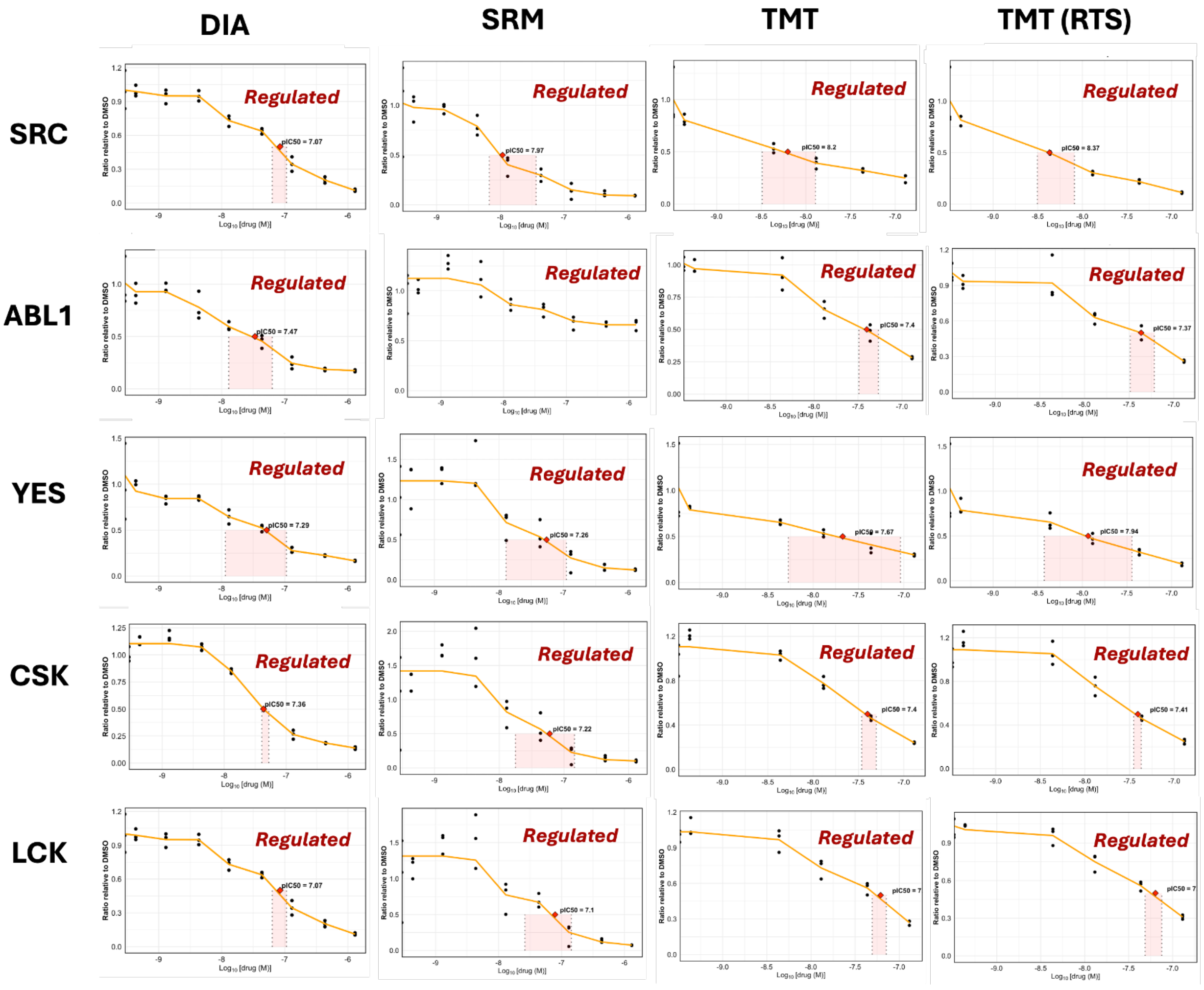

Figure 8: **Benchmark dataset: all positive controls exhibited a decreasing response upon increasing the dose of dasatinib.** Similar patterns were observed in all four benchmark datasets. In cases where 50% inhibition is not met (e.g., ABL1 in SRM benchmark), MSstatsResponse does not return OC50 estimates. In the TMT benchmark, YES was a borderline case with an adjusted p-value of 0.06 (cutoff = 0.05).

#### 4.3 In experimental datasets, statistical methods produced distinct dose-response curves

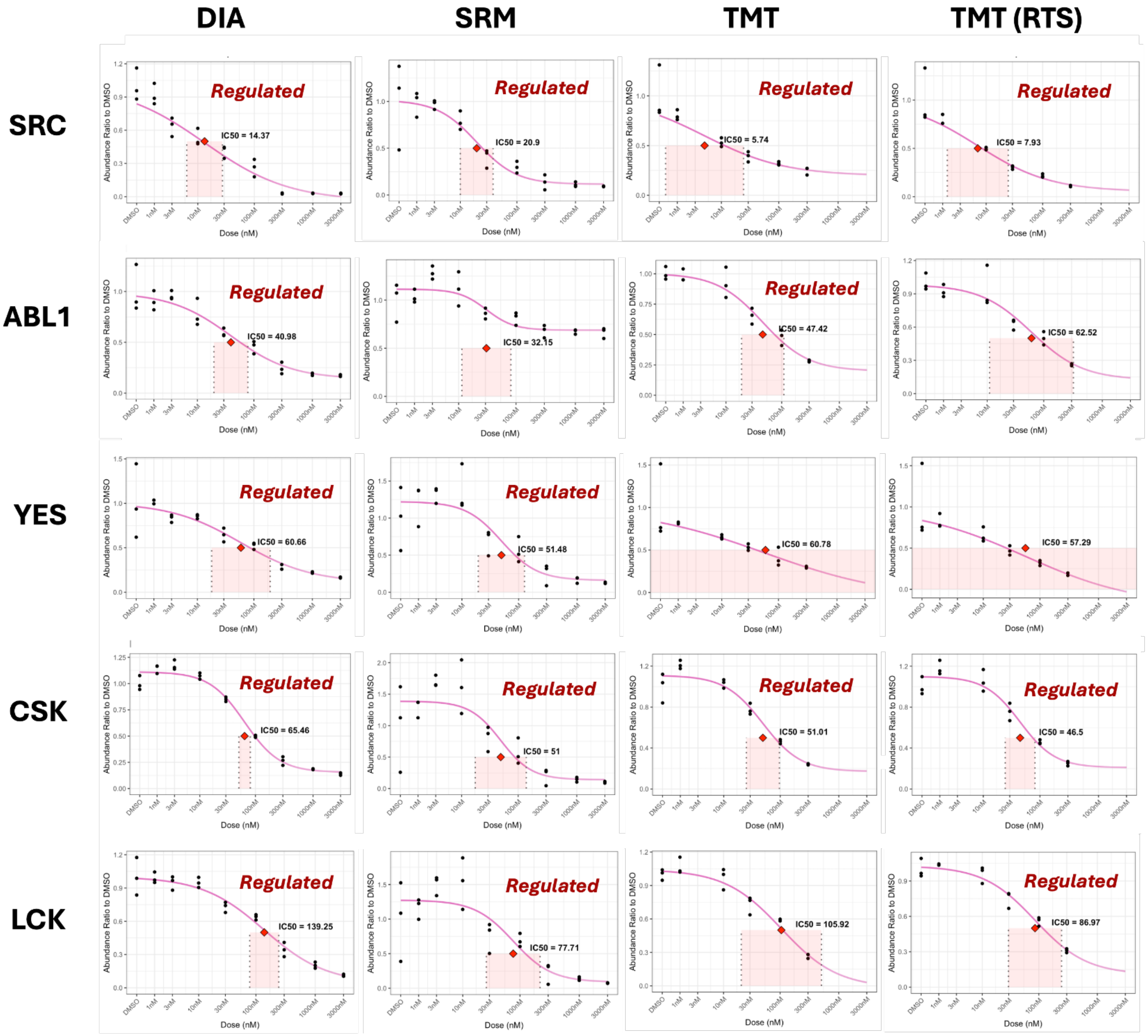

Figure 9: **Benchmark dataset: dr4pl response curves of all positive control proteins across acquisition benchmark datasets.** Similar patterns were observed in all four benchmark datasets. In cases where a sigmoidal trend was not observed (e.g., YES in the TMT and TMT-RTS benchmarks), dr4pl was unable to accurately estimate OC50 values, resulting in unbounded or large confidence intervals.

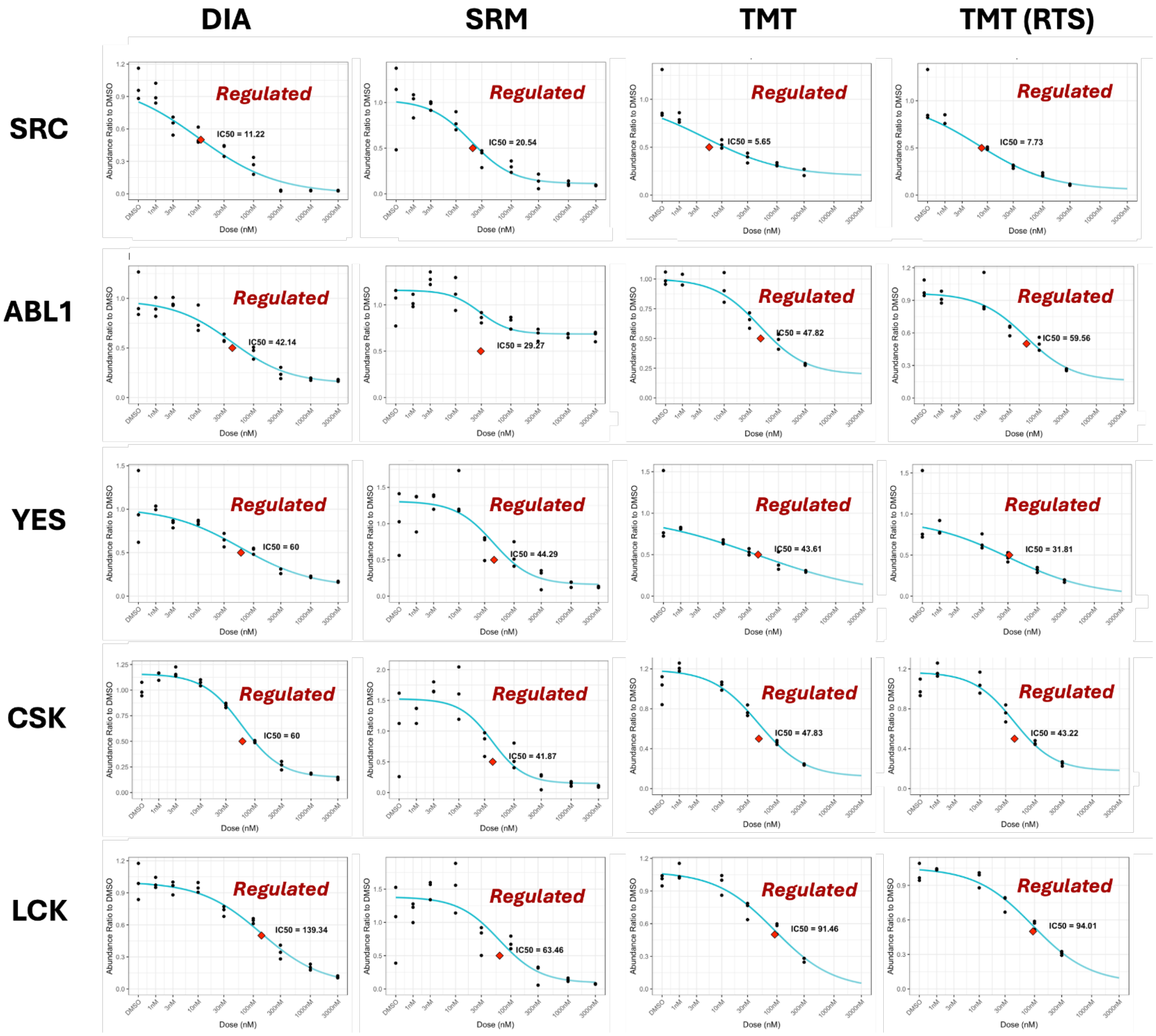

Figure 10: **Benchmark dataset: CurveCurator response curves of all positive control proteins across acquisition benchmark datasets.** Similar patterns were observed in all four benchmark datasets. In cases where partial inhibition is observed (e.g., ABL1 in SRM benchmark), CurveCurator failed to pick up the interaction.

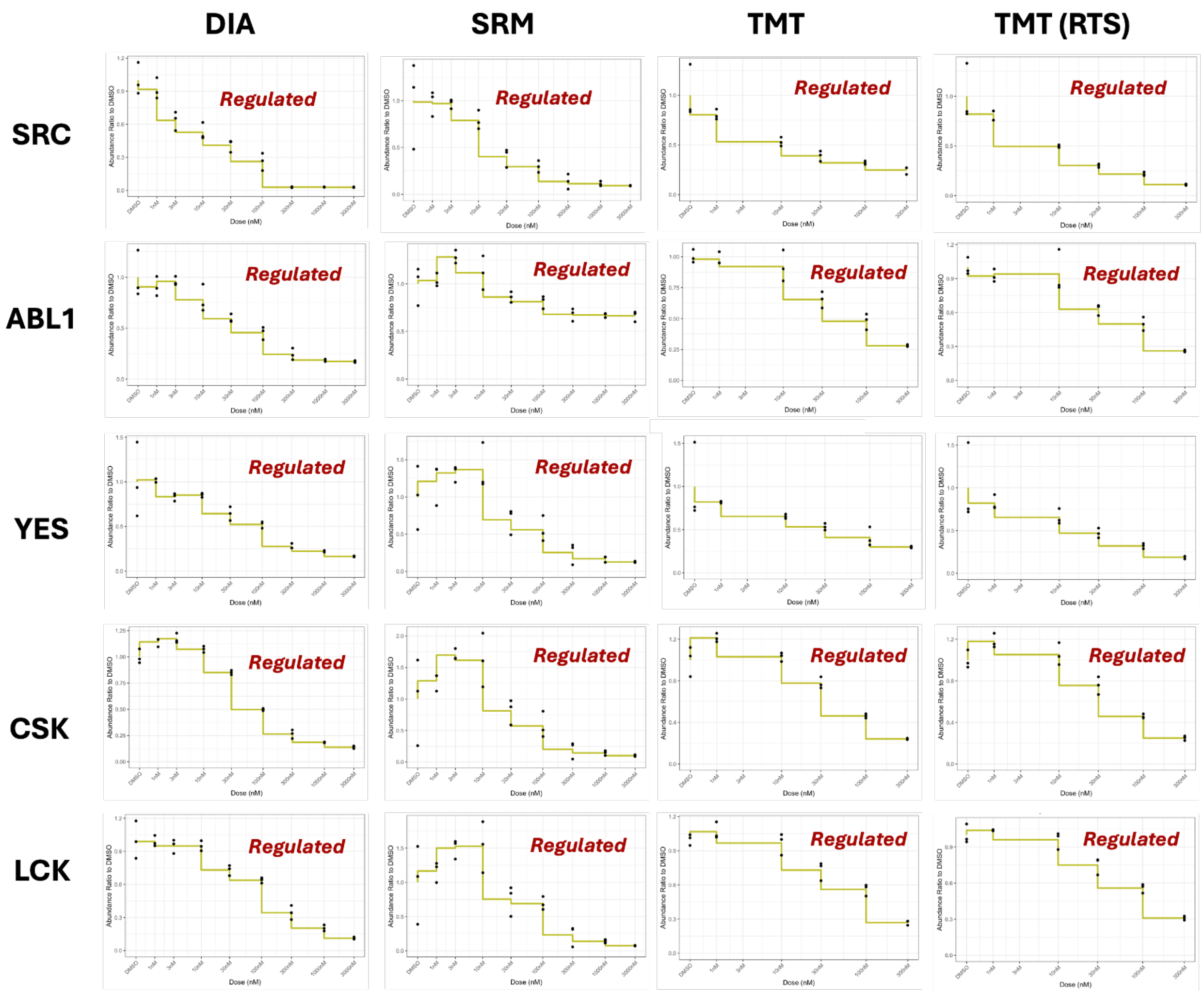

Figure 11: **Benchmark dataset: ANOVA response curves of all positive control proteins across acquisition benchmark datasets.** The ANOVA model fits to group means exactly, and is not well suited for does-response trend visualizations. Response patterns varied across proteins. For example, LCK and CSK in the SRM benchmark showed an initial increase followed by a decrease in abundance. ANOVA also failed to detect YES protein in both TMT benchmarks.

##### 4.4 In experimental datasets, statistical methods differed in drug-target detection sensitivity

| | TMT doses [0, 1, 10, 30, 100, 300] | | | | All doses | |
|---|---|---|---|---|---|---|
| Method | DIA | TMT | TMT w/RTS | SRM | DIA | SRM |
| drc | 0 | 5 (2) | 5 (1) | 0 | 5 (4) | 2 (0) |
| dr4pl | 2 (2) | 6 (3) | 7 (3) | 0 | 9 (5) | 4 (4) |
| CurveCurator | 13 (5) | 10 (5) | 12 (5) | 6 (4) | 26 (5) | 9 (4) |
| ANOVA | 10 (3) | 6 (4) | 10 (4) | 0 (0) | 8 (5) | 5 (5) |
| MSstatsResponse | 4 (4) | 9 (4) | 11 (5) | 1 (1) | 14 (5) | 5 (5) |

Table 2: **Benchmark dataset significant drug-protein interactions across acquisition types and methods.** Columns 1–4 report results using six drug doses (DMSO, 1nM, 10nM, 30nM, 100nM, 300 nM) across DIA, TMT, TMT with RTS, and Targeted datasets. Columns 5–6 expand to include all available doses for DIA and Targeted (including 3 nM, 1 µM, and 3 µM). The number of significant proteins is shown for each method. Known Dasatinib targets (SRC, ABL1, YES, CSK, and LCK) are shown in parentheses and serve as positive controls.

##### 4.5 In pseudo-single-replicate experimental datasets, drug-target detection varied between replicates

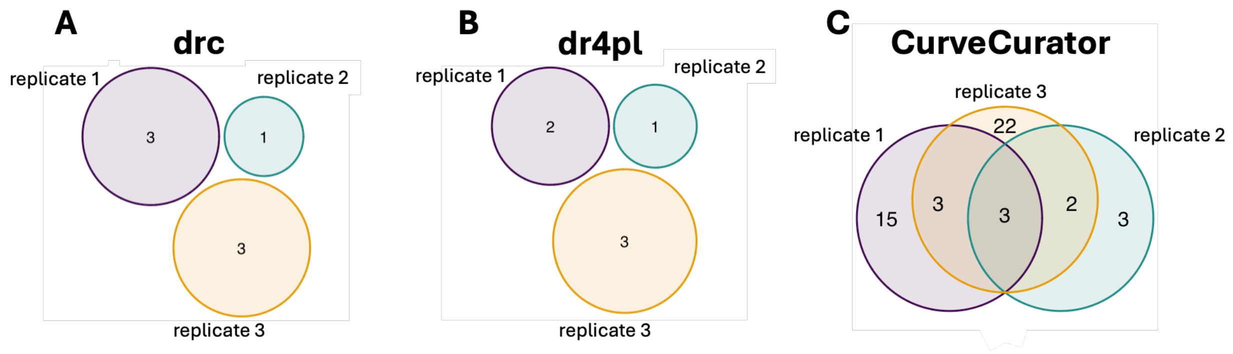

Figure 12: **TMT benchmark: single replicate analysis results vary greatly based on replicate choice.**

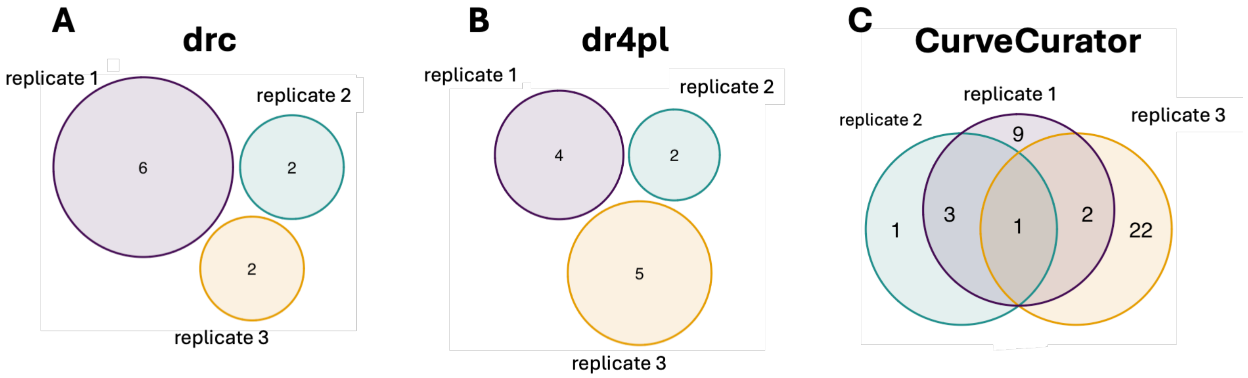

Figure 13: **TMT RTS benchmark: single replicate analysis results vary greatly based on replicate choice.**

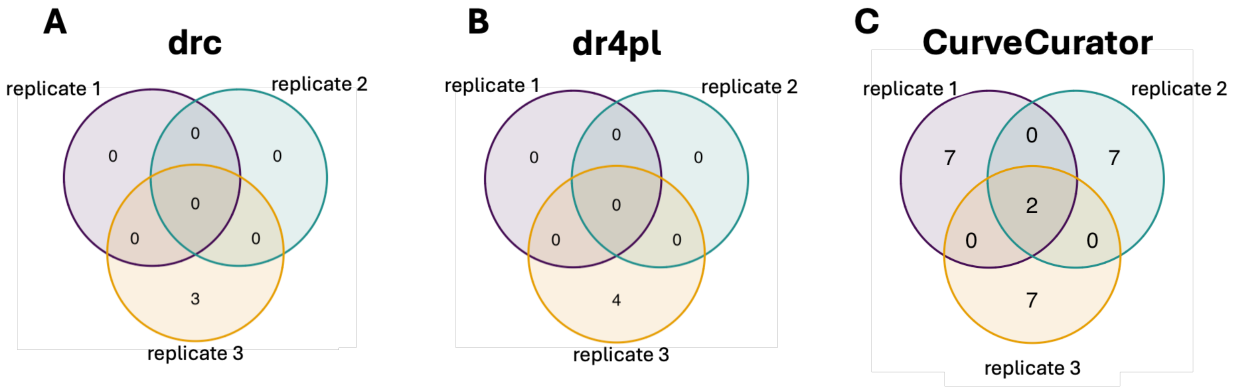

Figure 14: **SRM benchmark: single replicate analysis results vary greatly based on replicate choice.**

##### 4.6 In simulated datasets, MSstatsResponse detected weak interactions without increasing false positives

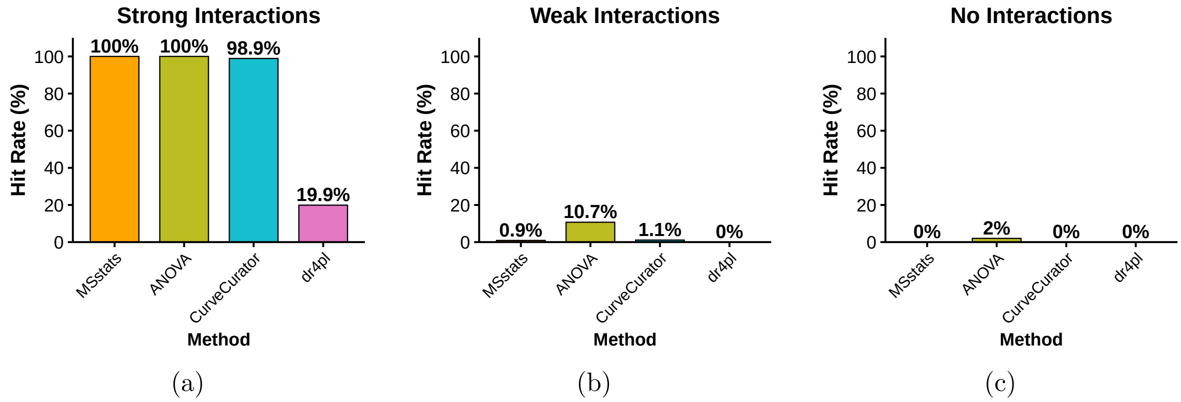

Figure 15: **Simulated dataset: method performance in high-variability confirmatory experiments.** Performance was evaluated on 3,000 simulated proteins across 9 doses (DMSO, 1nM, 3nM, 10nM, 30nM, 100nM, 300nM, 1000nM, 3000nM) with 3 replicates per dose and variance = 0.8. **(a)** Strong interactions: dr4pl failed to handle high variability, with significantly decreased hit detection, while MSstats, ANOVA, and CurveCurator remained stable. **(b)** Weak interactions: all methods failed to pick up most weak interactions here, only ANOVA detected a small amount of weak interactions. **(c)** Non-interactions: all methods correctly avoided false positives.

#### 4.7 In simulated datasets with fewer doses, MSstatsResponse maintained drug-target detection sensitivity

| Dose count | Method | Acquisition method: DIA | TMT | TMT-RTS | SRM |
|---|---|---|---|---|---|
| 3 doses | drc | 1 (0) | 0 | 1 (0) | 0 |
| | dr4pl | 0 | 0 | 0 | 0 |
| | CurveCurator | 24 (5) | 10 (5) | 11(5) | 5 (4) |
| | ANOVA | 12 (5) | 8 (4) | 16 (5) | 5 (4) |
| | MSstatsResponse | 6 (5) | 6 (4) | 10 (5) | 3 (3) |
| 1 dose | drc | 0 | 0 | 0 | 0 |
| | dr4pl | 0 | 0 | 0 | 0 |
| | CurveCurator | N/A | N/A | N/A | N/A |
| | ANOVA | 6 (3) | 3 (2) | 2 (1) | 0 |
| | MSstatsResponse | 2 (2) | 3 (2) | 2 (1) | 0 |

Table 3: **Benchmark dataset significant drug–protein interactions across acquisition types and dose subsets.** Results are shown for analyses using a DMSO control and either a three-dose series—1 nM, 1 µM, and 3 µM for DIA and SRM; and 1 nM, 100 nM, and 300 nM for TMT—or a single high-dose treatment—3 µM for DIA and SRM, and 300 nM for TMT. Columns report the number of significant proteins detected for each acquisition method and analysis approach. Known Dasatinib targets (SRC, ABL1, YES, CSK, and LCK) are shown in parentheses and serve as positive controls.
